# The sterol C-24 methyltransferase encoding gene, *erg6*, is essential for viability of *Aspergillus* species

**DOI:** 10.1101/2023.08.08.552489

**Authors:** Jinhong Xie, Jeffrey M. Rybak, Adela Martin-Vicente, Xabier Guruceaga, Harrison I. Thorn, Ashley V. Nywening, Wenbo Ge, Josie E. Parker, Steven L. Kelly, P. David Rogers, Jarrod R. Fortwendel

## Abstract

Ergosterol is a critical component of fungal plasma membranes. Although many currently available antifungal compounds target the ergosterol biosynthesis pathway for antifungal effect, current knowledge regarding ergosterol synthesis remains incomplete for filamentous fungal pathogens like *Aspergillus fumigatus*. Here, we show for the first time that the lipid droplet-associated sterol C-24 methyltransferase, Erg6, is essential for *A. fumigatus* viability. We further show that this essentiality extends to additional *Aspergillus* species, including *A. lentulus, A. terreus,* and *A. nidulans*. Neither the overexpression of a putative *erg6* paralog, *smt1,* nor the exogenous addition of ergosterol could rescue *erg6* deficiency. Importantly, Erg6 downregulation results in a dramatic decrease in ergosterol and accumulation in lanosterol and is further characterized by diminished sterol-rich plasma membrane domains (SRDs) at hyphal tips. Unexpectedly, *erg6* repressed strains demonstrate wild-type susceptibility against the ergosterol-active triazole and polyene antifungals. Finally, repressing *erg6* expression reduced fungal burden accumulation in a murine model of invasive aspergillosis. Taken together, our studies suggest that Erg6, which shows little homology to mammalian proteins, is potentially an attractive antifungal drug target for therapy of *Aspergillus* infections.

**IMPORTANCE:** *A. fumigatus* is the most common pathogen that causes invasive aspergillosis, a life-threatening fungal infection with more than 300,000 cases reported annually. Available antifungals to treat *Aspergillus*-related infection are limited to three drug classes targeting the plasma membrane (ergosterol) or the cell wall, each of which suffer from either host toxicity or rising resistance levels. As ergosta-type sterols are absent in mammalian cells but are essential for fungal viability, the ergosterol biosynthesis pathway remains an enticing target for the development of new antifungals. Although ergosterol biosynthesis has been well studied in model yeast, only a few genes have been genetically characterized in *A. fumigatus*. Here, we characterize Erg6, one of the fungus-specific sterol biosynthesis genes, as an essential gene in *Aspergillus* species. We further provide *in vivo* evidence of the importance of Erg6 for establishment of invasive aspergillosis. Given the importance of Erg6 in other fungal systems for growth, stress resistance, and virulence, our study suggests that development of Erg6 inhibitors may be a promising strategy for developing novel broad-spectrum antifungals.

## INTRODUCTION

*Aspergillus fumi*gatus is the most prevalent *Aspergillus* species that causes invasive Aspergillosis (IA), a life-threatening fungal infection with high mortality rates up to 40% - 50% (1). With the increasing numbers of patients having immune defects, *Aspergillus*-related infections have become an important public health concern (2). Currently, there are only three available classes of antifungal compounds for the treatment of IA (i.e., triazoles, polyenes, and echinocandins), all targeting essential components of the fungal cell membrane or cell wall (3). Unfortunately, the clinical efficacy of these antifungal classes is hampered by host toxicity, side effects, and poor bioavailability to some extent. Moreover, the global emergence of resistance, especially to the triazole class, makes their clinical application for long-term treatment more complicated (2, 4).

Sterols are functional and constructional components residing in the plasma membrane and are responsible for the cell membrane permeability, fluidity, and stability (5). Ergosterol (C28 sterol) is a specific sterol found in fungi, plants, and protozoa, whereas mammalian cells synthesize cholesterol (C27 sterol) as the major membrane sterol (6). Because of the uniqueness and essentiality of ergosterol for fungal organisms, disturbing fungal ergosterol homeostasis is widely considered a promising strategy for novel antifungal development (7). Besides triazoles and polyenes, statins and allylamines are two classes of inhibitors targeting enzymatic steps in the ergosterol biosynthesis pathway (8, 9). Ergosterol and cholesterol are both sterols with similar four-ring structure harboring a hydroxyl group at C-3 and an unsaturated bond at C-5,6. The distinguishing feature between ergosta-type and cholesta-type sterols is that ergosta-type sterols contain a methyl group at C-24 on the side chain. Addition of this methyl group is catalyzed by the sterol C-24 methyltransferase enzyme, encoded by the *erg6* gene in fungi (10). As fungi and humans are eukaryotic organisms, most enzymes involved in fungal ergosterol biosynthesis have homologs in mammalian cholesterol biosynthesis. However, Erg6 is one of the three unique enzymes that is absent in the human cholesterol biosynthesis pathway (11).

Among the organisms studied to date, yeast and filamentous fungi appear to share conserved early and late enzymatic steps of the ergosterol biosynthesis pathway. However, after the formation of the first sterol-type intermediate, lanosterol, the pathway bifurcates into one of two paths. In budding yeast, like *S. cerevisiae*, lanosterol is catalyzed to 4,4-dimethylcholesta-8,14,24-trienol by the triazole-target gene, Erg11 (5). As for *A. fumigatus*, eburicol is the preferred substrate of the Erg11 orthologs, Cyp51A and Cyp51B, and eburicol is generated from lanosterol by the activity of Erg6 (12). Therefore, although Erg6 represent one of the late enzymatic steps for ergosterol biosynthesis in *S. cerevisiae*, its substrate specificity for lanosterol makes it an early enzymatic step for organisms like *A. fumigatus*. Erg6 catalyzes a methyl addition to C-24 by the way of an S-adenosylmethionine (SAM)–dependent transmethylation and shifts a double bond to produce a C-24(28)-methylene structure with high substrate specificity (13). Erg6 has been genetically characterized in multiple single-celled yeast, including *S. cerevisiae*, *Kluyveromyces lactis*, *Candida glabrata*, *Candida albicans*, *Cryptococcus neoformans*, and *Pneumocystis carinii* (14–18). Deletion of *erg6* is not lethal for these fungi, however, the *erg6* loss-of-function mutations cause alternation in drug susceptibility and defective growth phenotypes related to membrane integrity and permeability. Studies of Erg6 for filamentous fungi are relatively limited. Recently, Erg6 has been characterized in *Mucor lusitanicus*. Unlike yeast genomes that encode only one copy of *erg6*, *M. lusitanicus* possesses three copies of Erg6, referred to Erg6A, Erg6B, and Erg6C (19). Erg6B plays a critical role in ergosterol biosynthesis. Deletion of *erg6B* compromises ergosterol production, growth ability, antifungal resistance and virulence, and double deletion together with *erg6A* or *erg6C* is lethal for *M. lusitanicus* (19).

In this study, we characterized Erg6 in *A. fumigatus*. Although *A. fumigatus* encodes two putative sterol C-24 methyltransferase, designated as Erg6 and Smt1, we report that only loss of *erg6* generates significant phenotypes. Strikingly, we find that *erg6* is essential for *A. fumigatus* viability *in vitro* and for disease establishment in a murine model of invasive aspergillosis. We also show that *erg6* orthologs are essential across multiple *Aspergillus* species. Repression of *A. fumigatus erg6* expression in a conditional mutant blocked ergosterol biosynthesis resulting in abundant accumulation of lanosterol, the proposed substrate of Erg6. Surprisingly, the downregulation of *erg6* did not drive significant changes in triazole or polyene susceptibility profiles. This result is contrary to *erg6* mutants in other fungal species. Taken together, our data support inactivation of Erg6 as a promising therapeutic approach for fungal infection.

## RESUTLS

### Erg6 is indispensable for *A. fumigatus* viability

To identify putative *A. fumigatus* orthologs of *S. cerevisiae ERG6*, we performed a BLASTP analysis using the amino acid sequence of *S. cerevisiae* Erg6p (SGD: S000004467) against the *A. fumigatus* genome database (fungidb.org). Two putative protein-encoding loci, AFUB_099400 (EDP47339, 54.18% identity) and AFUB_066290 (EDP50296, 31.39% identity), which are designated as *erg6* and *smt1*, respectively, were identified. Alignment analysis showed that the putative *A. fumigatus* Erg6 and Smt1 proteins share 26.71% amino acid identity with each other. To explore the phylogenetic relationship of sterol C-24 methyltransferase in different fungi, the same analysis was performed in *A. lentulus*, *A. terreus*, *A. nidulans*, *S. cerevisiae*, *C. albicans*, *C. neoformans*, and *Neurospora crassa*. A phylogenetic tree was constructed based on full-length amino acid sequences using the maximum likelihood method (Fig. S1). Remarkably, only a single sterol C-24 methyltransferase encoding gene was found in the yeast organisms analyzed, whereas the filamentous fungi analyzed each harbored at least two putative paralogs (Fig. S1).

To investigate the importance of sterol C-24 methyltransferase activity in *A. fumigatus*, we first attempted to generate null mutants of both *erg6* and *smt1*, completely replacing the open reading frames (ORFs) with a hygromycin selection cassette, using a highly efficient CRISPR/Cas-9 gene editing technique (Fig. S2A) (20). No *Δerg6* transformant was obtained after several transformation attempts, whereas *Δsmt1* mutants were successfully generated. These results implied a differential requirement for the two putative sterol C-24 methyltransferases in *A. fumigatus*, with the *erg6* homolog potentially being essential. Phenotypic analysis of the Δ*smt1* mutant revealed no differences in colony growth or morphology when compared to the control strain, suggesting that either Smt1 plays a minimal role in ergosterol biosynthesis or that Erg6 activity is able to compensate for loss of *smt1* (Fig. S3A). To generate a hypomorphic allele of *erg6* for further study, we next constructed a pTetOff*-erg6* mutant in which the endogenous *erg6* promoter was replaced by a tetracycline-repressible promoter (Fig. S2B) (21). Although this genetic manipulation resulted in a 4-fold (log_2_) increase in *erg6* expression in the absence of doxycycline, the presence of only 0.5 µg / ml doxycycline in the culture media generated a 4-fold (log_2_) reduction in gene expression (Fig. 1A). Importantly, the pTetOff*-erg6* mutant displayed growth and colony morphology identical to the parental strain when cultured in the absence of doxycycline, suggesting that the basal upregulation of *erg6* expression in the absence of doxycycline does not alter basic growth of *A. fumigatus* (Fig. 1B, left panel). However, growth and germination were significantly inhibited in the pTetOff-*erg6* strain in the presence of increasing doxycycline concentrations (Fig. 1B and 1C). As low as 0.5 μg / ml doxycycline almost completely prevented colony development on solid agar (Fig. 1B). Analysis of submerged culture demonstrated that the pTetOff*-erg6* strain exhibited a dose-dependent decrease in mycelial development in response to increasing doxycycline concentrations (Fig. 1C).

**Figure 1.**
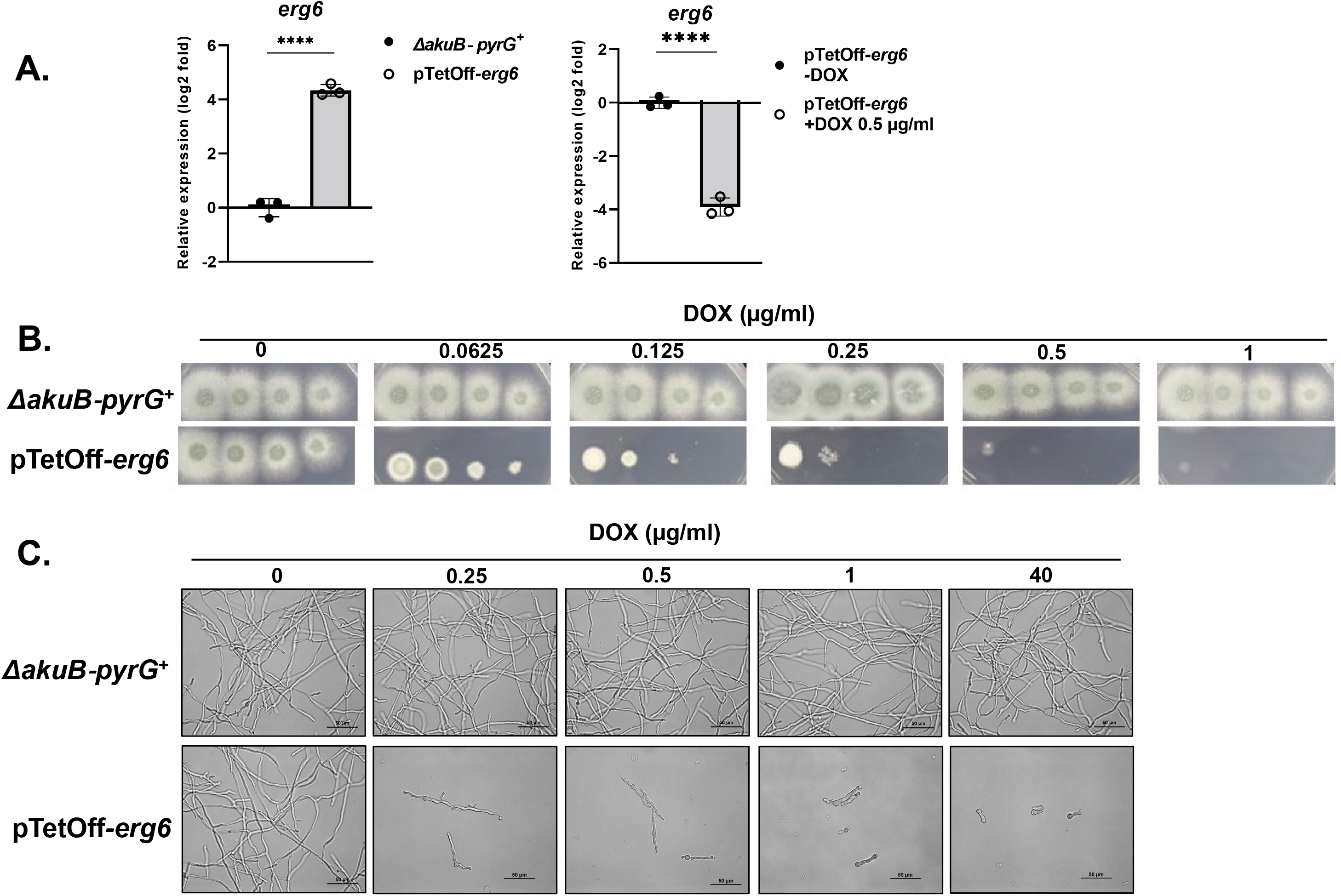
Repression of *erg6* expression inhibits *A. fumigatus* growth *in vitro*. **(A)** The expression level of *erg6* in indicated conditions as analyzed by RT-qPCR. Mycelia were harvested after 16 h in liquid GMM at 37°C, 250 rpm. Gene expression was normalized to the reference gene, *tubA*, and data presented as mean ± SD of log_2_ fold change. All assays were performed in biological triplicate. Two-tailed Student t-test was used for statistical analysis. ***p<0.0005, ****p<0.0001. **(B)** Spot-dilution assays were performed on GMM agar plates with the parental and pTetOff-*erg6* strains in the indicated doxycycline levels. For all assays, suspension aliquots of 5 μl containing 50,000, 5,000, 500, and 50 total conidia were inoculated and plates were incubated at 37°C for 48 h. **(C)** Microscopic images of the parental and pTetOff-*erg6* strains after 16 h of exposure to the indicated doxycycline levels in static GMM culture at 37 °C. Microscopy was performed on Nikon NiU with bright field settings.

Although these findings suggested that *erg6* is likely essential for *A. fumigatus* viability, the pTetOff-*erg6* conidia were able to germinate and establish initial polarity in the presence of doxycycline, even as high as 40 µg / ml (Fig. 1C). Therefore, Erg6 activity could potentially only be essential for growth and viability post-polarity establishment. To further test if *erg6* is differentially essential for pre- or post-germination viability, we next constructed a pTetOn-*erg6* mutant (Fig. S2B) that should require the presence of doxycycline for *erg6* expression (22). Importantly, the pTetOn-*erg6* mutant behaved as expected in culture, with colony development only occurring upon addition of doxycycline to the media (Fig. S4). Both pTetOn-*erg6* and pTetOff-*erg6* stains were employed in live-cell staining assays using the fluorescent marker 5-carboxyfluorescein diacetate (CFDA) (23). Parental and pTetOn-*erg6* conidia were cultured in GMM broth with or without 100 µg / ml doxycycline for 16 hours, followed by CFDA staining to quantitatively measure viability of germlings. Germlings that were either fully or only partly CFDA-labeled were counted as viable cells. The cultures were limited to 16 hours of incubation to allow unambiguous detection of live *vs*. dead (i.e., stained *vs*. unstained) fungal elements. As shown in Fig. 2B, upper panel, without doxycycline as an inducer, no polarity was observed in pTetOn-*erg6* with rare viable conidia stained with CFDA. With 100 µg / ml doxycycline, the pTetOn-*erg6* mutants were induced and germinated with a comparable viability rate with background strains (Fig. 2A, left panel). Thus, these findings confirm that *erg6* is required for *A. fumigatus* viability beginning with the earliest stages of growth.

**Figure 2.**
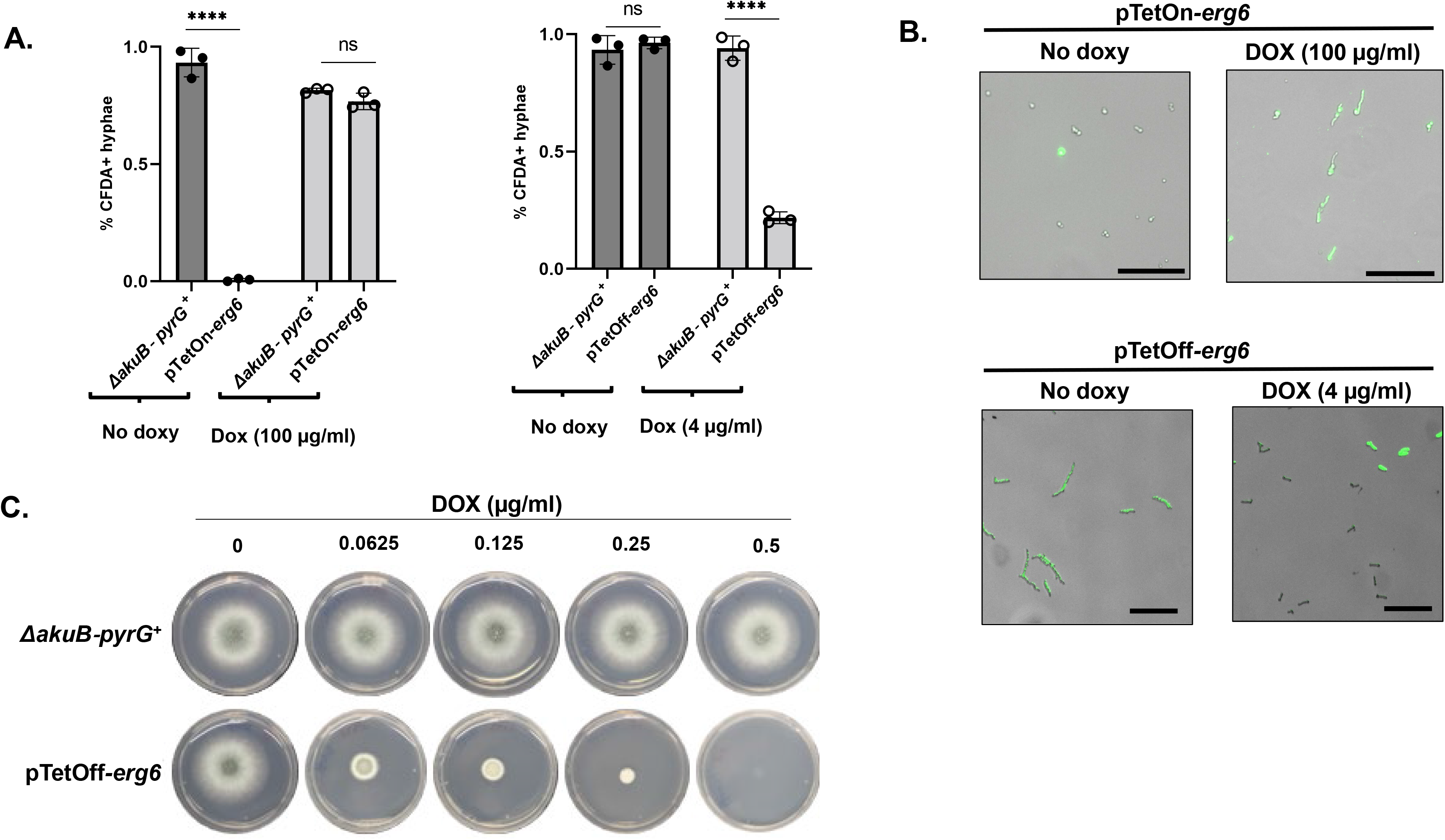
Erg6 is essential for *A. fumigatus* viability. Viability rates of pTetOn-*erg6* **(A)** and pTetOff-*erg6* **(B)** in the indicated doxycycline levels using a CFDA staining assay. Hyphae were harvested after GMM culture for 16 h at 30°C and subsequently stained with 50 μg/ml CFDA for 1 h. Microcolonies that showed bright a green signal were manually enumerated as viable. More than 200 microcolonies were measured in each assay and all experiments were completed in triplicate. Data is depicted as the mean ± SEM. Two-tailed Student t-tests were used for statistical analysis. ****p<0.0001. **(C)** Conidia of the parental and pTetOff-*erg6* strains were initially grown in GMM broth without doxycycline treatment for 8 h to allow formation of germlings. Subsequently, germling aliquots of 10 µl were transferred to fresh GMM agar plates supplemented with the indicated concentration of doxycycline, and plates were incubated for an additional 48 h.

Similar results were achieved when using the pTetOff-*erg6* mutant. Parental and pTetOff-*erg6* conidia were cultured in GMM broth with or without 4 µg / ml doxycycline for 16 hours. Nearly all cells for parental and pTetoff-*erg6* strains cultured in doxycycline-free conditions were positively fluorescently labeled with CFDA, indicating 100% viability in non-repressive conditions. In the presence of 4 µg / ml doxycycline, whereas the parental strain remained unaffected, the pTetoff-*erg6* displayed a sharp decrease in the CFDA-labeled population of germlings with only 20% positivity (Fig. 2A, right panel). To further confirm *erg6* deficiency is lethal for *A. fumigatus*, conidia of parental and pTetoff-*erg6* strains were cultured to the germling stage in the GMM broth without doxycycline and subsequently inoculated onto GMM agar containing increasing concentrations of doxycycline. As shown in Fig. 2C, similar to the hyphal growth inhibition exhibited by culturing conidia on doxycycline-impregnated agar plates, pre-formed germlings of the pTetOff-*erg6* mutant were entirely growth inhibited with as little as 0.5 µg / ml doxycycline. Taken together, these findings demonstrate that *erg6* deficiency is lethal for *A. fumigatus*.

To determine if the essentiality of *erg6* was not only specific to *A. fumigatus* but may instead be generalizable across *Aspergillus* species, we next generated tetracycline-repressible promoter replacement mutants targeting the *erg6* orthologs of *A. lentulus, A. terreus,* and *A. nidulans* through the CRISPR/Cas-9 technique. Erg6 orthologs were retrieved from the most similar alignments in BLASTP analysis against the genome databases of the respective *Aspergillus* species using the amino acid sequence of *S.* cerevisiae *erg6* (SGD: S000004467) as a query sequence. As we noted for *A. fumigatus*, there was a clear negative correlation between increasing doxycycline concentrations and colony establishment on agar plates among three additional filamentous *Aspergillus* species (Fig. 3). Thus, *erg6* is essential in multiple *Aspergillus* species.

**Figure 3.**
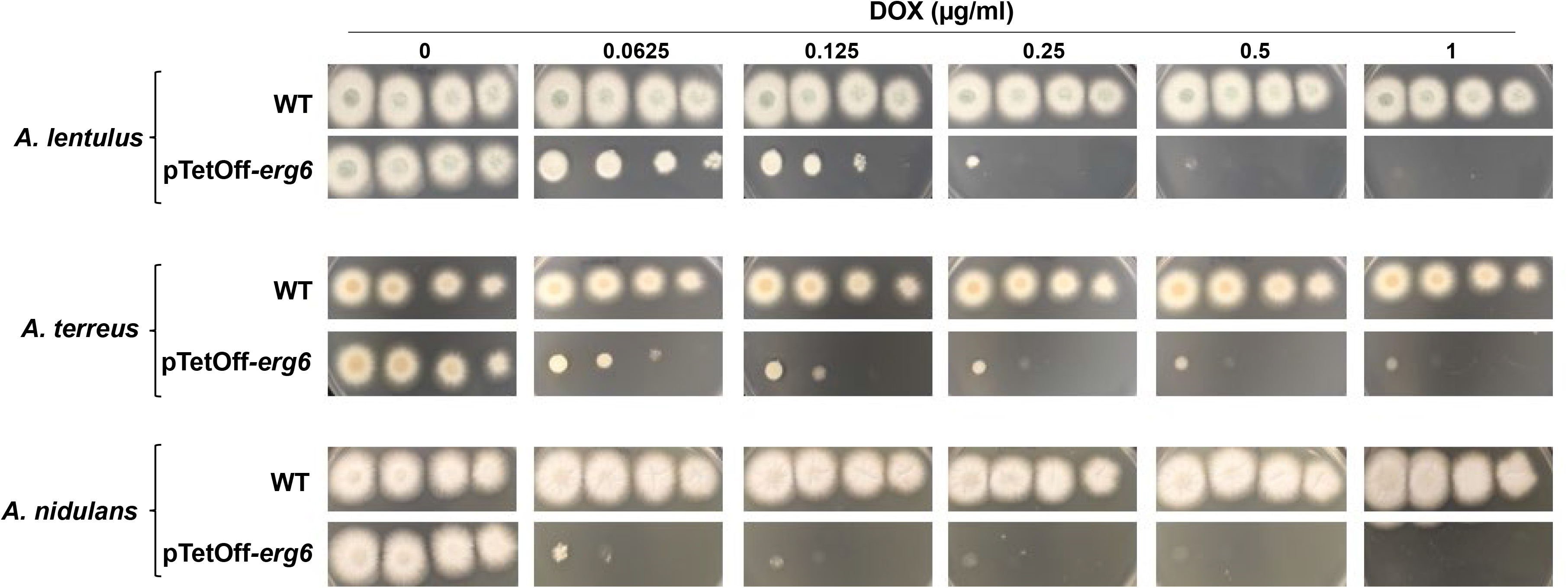
Erg6 is essential across *Aspergillus* species. Colony morphology of parental and pTetOff-*erg6* strains of *A. lentulus*, *A. terreus*, and *A. nidulans* in the presence of the indicated concentrations of doxycycline. Spot-dilution culture was performed as described in Figure 1B. For *A. lentulus* and *A. terreus*, conidia were inoculated onto GMM plates, whereas *A. nidulans* conidia were cultured on GMM supplemented with 5% yeast extract.

### Overexpression of *smt1* or exogenous sterols cannot rescue loss of *erg6*

We next sought to examine the functional relationship between *erg6* and the predicted paralog, *smt1*. To address whether overexpression of *smt1* could rescue *erg6* repression, we constructed a pTetOff-*erg6* mutation in a *smt1* overexpression background (Fig. S2B). The *smt1* endogenous promoter was first replaced with the strong p*HspA* promoter (24) through CRIPSR/Cas9-mediated gene targeting to generate strain OE-*smt1* (Fig. S2B). Although this promoter replacement generated a ~6-fold (log2) upregulation of *smt1* expression, growth and colony development were unaffected (Fig. S3A and S3B). The pTetOff-*erg6* promoter construct was then integrated in the OE-*smt1* genetic background. When these mutants were employed in spot-dilution assays, constitutive *smt1* overexpression driven by the p*HspA* promoter was not able to promote colony development when *erg6* was downregulated by doxycycline addition (Fig. 4A). Further, using RT-qPCR to measure *erg6* and *smt1* expression levels, we found that the expression of neither *erg6* nor *smt1* was responsive to loss of the other (Fig. S3C). To further rule out the possibility that *smt1* compensates for *erg6* deficiency, a pTetOff-*erg6* mutation was constructed in the *Δsmt1* genetic background. As shown in Fig. 4B, deletion of *smt1* was not found to exacerbate loss of viability when *erg6* expression was repressed by addition of exogenous doxycycline. An almost complete lack of colony development was evident at 0.5 µg / ml doxycycline as was seen in pTetOff-*erg6* strains expressing *smt1* (Figure 1C). Therefore, *smt1* does not appear to be a functional paralog of *erg6*.

**Figure 4.**
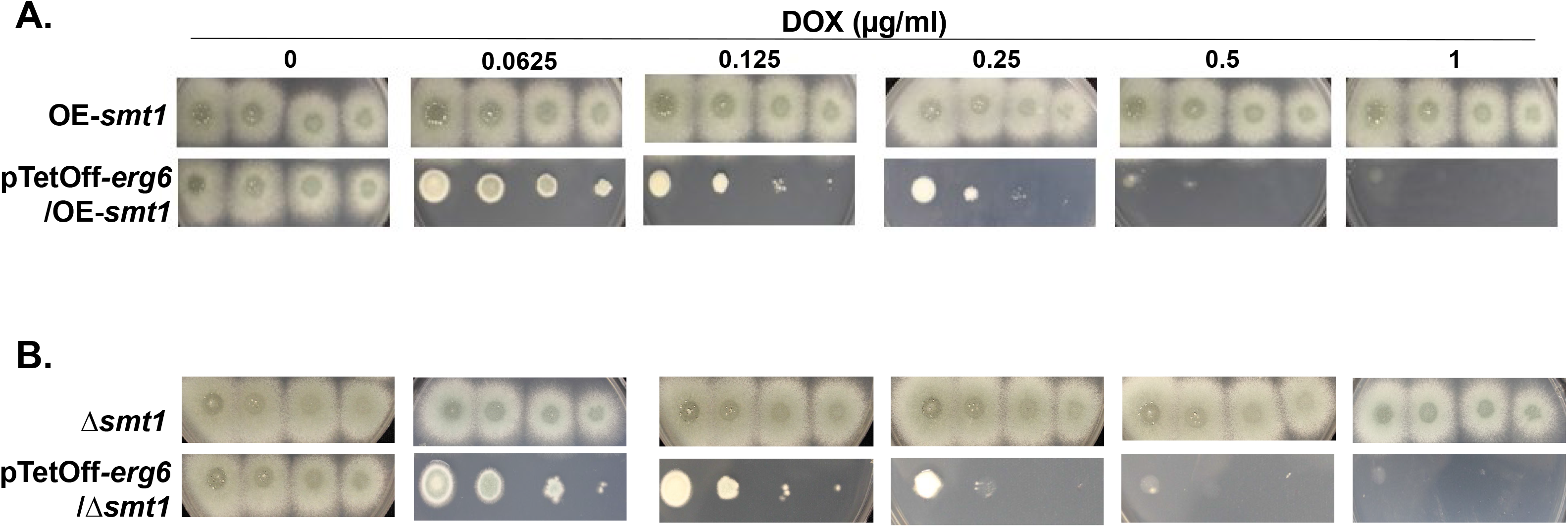
The putative paralog, *smt1*, shows no functional redundancy with *erg6*. Spot-dilution assays of pTetOff-*erg6* mutants constructed in the OE-*smt1* **(A)** or Δ*smt1* **(B)** genetic backgrounds. Culture conditions were as described in Figure 1B.

Previous studies indicated that *A. fumigatus* can uptake exogenous sterol to compensate for the stress induced by triazole ergosterol biosynthesis inhibitors, causing altered minimum inhibitory concentration endpoints (25). To see if exogenous sterols could at least partially suppress growth inhibition under doxycycline-mediated *erg6* repression, we performed similar growth assays on media embedded with either 10% fetal bovine serum (as a source of cholesterol) or ergosterol. Notably, we found that exogenous addition of neither ergosterol nor fetal bovine serum containing cholesterol were able to restore growth under *erg6* repression (Fig. 5). These findings indicate that exogenous sterols are unable to compensate for loss of viability incurred by *erg6* repression and suggest that our *in vitro* viability reduction may translate to the *in vivo* infection environment where potential cholesterol uptake could mask effects of ergosterol biosynthesis blockade.

**Figure 5.**
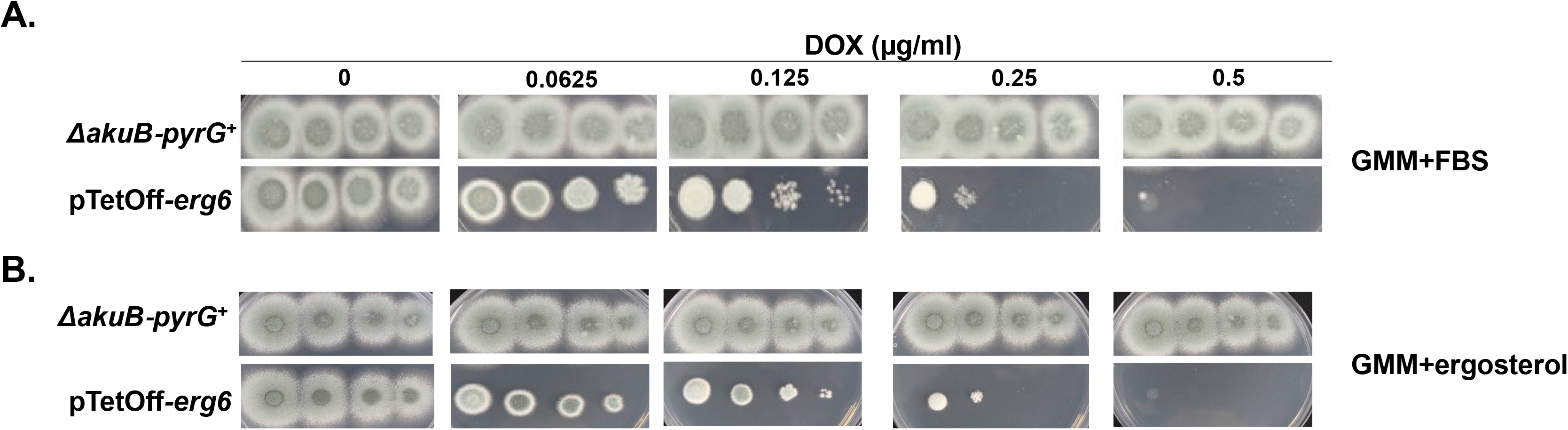
Exogenous supplementation with sterols does not restore growth due to loss of *erg6* expression. Spot-dilution assays of the parental and pTetOff-*erg6* strains were performed on GMM supplemented with 10% fetal bovine serum **(A)** or GMM supplemented with 40 μM ergosterol **(B)** at the indicated doxycycline concentrations. Culture conditions were as described in Figure 1B.

### Repression of *erg6* results in loss of membrane ergosterol and altered sterol profiles

Although *smt1* appeared to play little-to-no role in ergosterol biosynthesis to support growth, we next sought to ensure a conserved role for *erg6* in this important pathway for *A. fumigatus*. In filamentous fungi, ergosterol is known to accumulate in hyphal tips in structures called sterol-rich plasma membrane domains (SRDs), which have been validated to be essential cellular machinery involved in maintenance of growth polarity (26). Because *erg6* is a putative ergosterol biosynthesis pathway component and loss of *erg6* expression causes severe hyphal growth impairment, we hypothesized that *erg6* downregulation would result in the loss of ergosterol accumulation at hyphal tips, loss of total cellular ergosterol, and accumulation of the putative Erg6 substrate, lanosterol (12). To test this, we first stained hyphae of the parental and pTetOff-*erg6* strains with the sterol dye, filipin, which is widely used in filamentous fungi to visualize SRDs (26). As shown in Fig. 6A and 6B, filipin staining of the parental strain revealed concentrated fluorescence at the hyphal tips with and without doxycycline treatment, forming a cap-like pattern structure as indicated by the white arrows. In the absence of doxycycline, the pTetOff-*erg6* mutant behaved similarly (Fig. 6A, right panel). However, in the pTetoff-*erg6* doxycycline-treated cultures, the filipin staining pattern was completely disrupted with diminished hyphal staining and a loss of specific hyphal tip accumulation (Fig. 6B, right panel). To measure sterol profiles quantitatively, total sterols were harvested in trimethylsilane and analyzed using gas chromatography (GC)-mass spectrometry (MS) in both strains under increasing doxycycline concentrations. Sterol profiles of the parental strain were comparable in the presence or absence of doxycycline treatment, with ergosterol accounting for nearly 90% of total 24-methylated sterol and the Erg6 substrate, lanosterol, only accounting for ~0.6% (Table 1). As expected, the pTetOff-*erg6* mutant displayed sterol profiles similar to the parent strain when no doxycycline was added to the culture medium (Table 1). In contrast, among the pTetoff-*erg6* doxycycline treatment groups, the total ergosterol content decreased by almost 50% and lanosterol accumulated significantly to the second-most abundant measured sterol to nearly 40% of the total 24-methylated sterols present. Although lanosterol increased upon *erg6* repression in the pTetOff-*erg6* mutant, the Erg6 product, eburicol, remained relatively stable in the presence or absence of doxycycline with levels ranging from 0.9 - 1.0% for the parental strain and 1.3 - 2.4% for the pTetOff-*erg6* strain. In addition, several cholesta-type intermediates, including cholesta-5,7,22,24-tetraenol, cholesta-5,7,24-trienol, 4,4-dimethyl cholesta-dienol and cholesta-dienol, accounted for less than 4% of total 24-methylated sterols respectively in the doxycycline-treated pTetOff-*erg6* mutant, whereas these sterol intermediates were not detectable in the background strain. Taken together, these findings further confirm a conserved role for *A. fumigatus erg6* in ergosterol biosynthesis specifically at the lanosterol-to-ergosterol conversion step.

**Figure 6.**
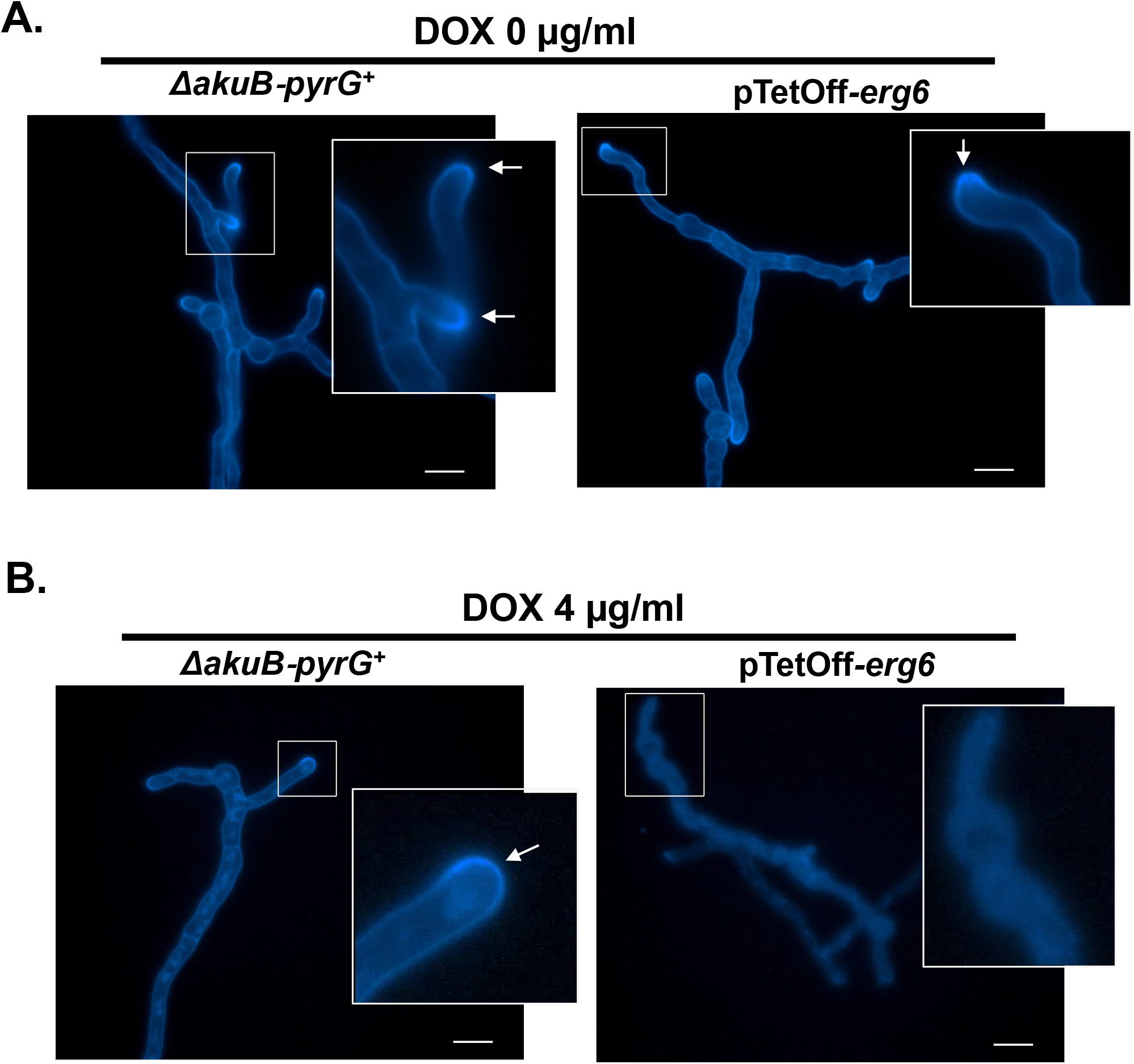
Repression of Erg6 expression alters ergosterol distribution in *A. fumigatus* hyphae. Mycelia of the parental and pTetOff-*erg6* strains were grown in GMM broth without doxycycline **(A)** and with 4 μg/ml doxycycline **(B)** for 16 h at 30°C. Hyphae were subsequently stained with 25 μg/ml filipin for 5 min. Fluorescent images were captured using DAPI filter settings. White arrows indicate sterol-rich plasma membrane domains (SRDs). Scale bar=10 µm.

**Table 1:**
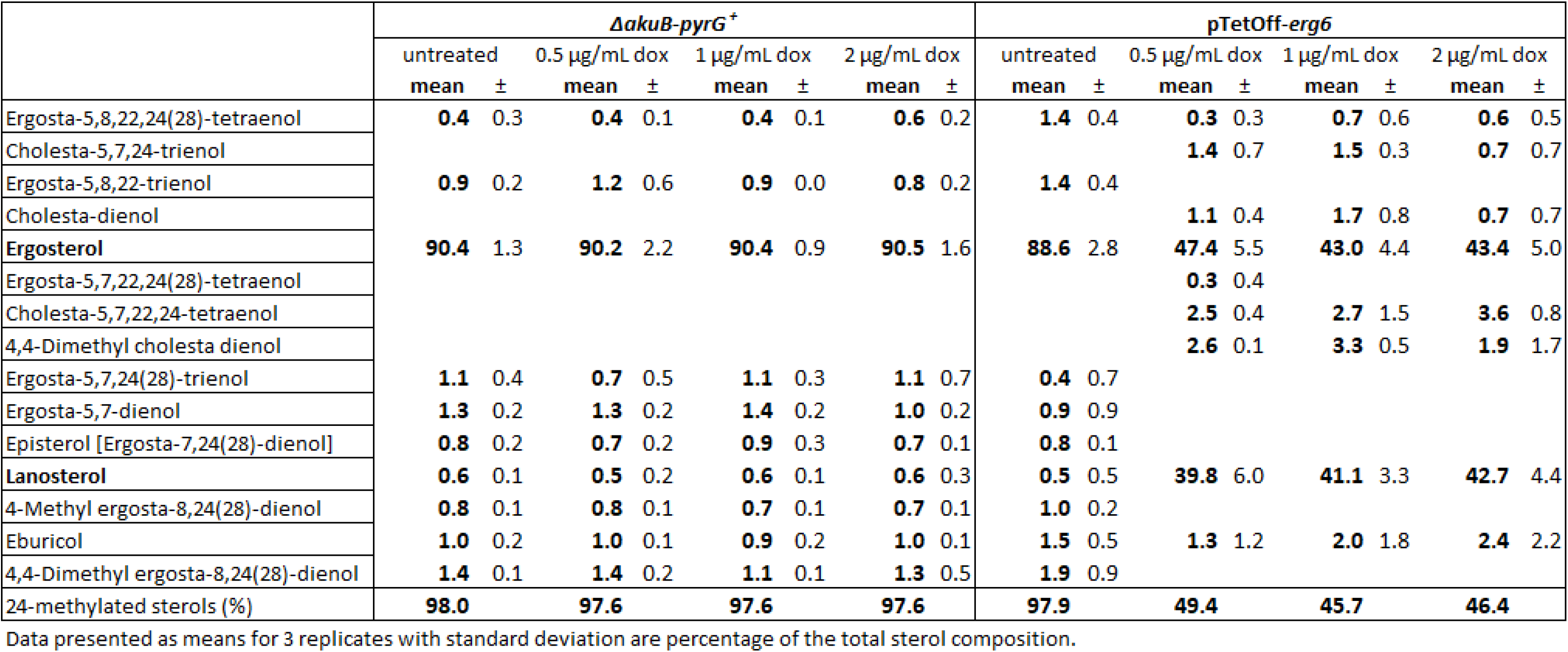
Changes in total 24-methylated sterol composition in response to *erg6* repression.

### *A. fumigatus* Erg6 localizes to lipid bodies

The Erg6 sterol C-24 methyltransferase homolog has been reported to localize to lipid droplets and to the endoplasmic reticulum of the model yeast, *S. cerevisiae* (27, 28). To examine if the localization of Erg6 in a filamentous pathogenic mold, like *A. fumigatus*, is conserved, we performed localization studies using strains expressing an Erg6-enchanced Green Fluorescent Protein (eGFP) chimera. Employing both the parental and pTetOff-*erg6* backgrounds, a construct was designed to fuse *egfp* to the 3’ end of *erg6*, such that *erg6-gfp* expression would be controlled by the native *erg6* promoter in the parental background and by the TetOff promoter in the pTetOff-*erg6* background (Fig. S2C). Phenotypic assays indicated that the Erg6-GFP mutants were functionally normal and that the pTetOff-*erg6-gfp* strain was as equally responsive as the non-chimeric mutant to doxycycline-mediated *erg6* repression (Fig. 7A). These results indicated that the GFP-fusion had no detrimental effects on Erg6. Fluorescent microscopic observation revealed that, regardless of doxycycline presence, Erg6-GFP displayed a punctate localization pattern distributed evenly throughout the mycelia of the *erg6-gfp* strain (Fig. 7B, upper panels). Notably, we observed that the Erg6-GFP signal in pTetOff-*erg6-gfp* strain cultured without doxycycline was much stronger than that of the *erg6-gfp* strain (Fig. 7B, lower left panel). These protein-level findings are consistent with our previous data showing basal overexpression of *erg6* when under the control of the pTetOff promoter and cultured in the absence of doxycycline (Fig. 1A, left panel). Regardless of protein abundance, Erg6-GFP localization remained confined to punctate structures dispersed throughout hyphae of the pTetOff-*erg6-gfp* strain in the absence of doxycycline. In contrast, the Erg6-GFP signal of the pTetOff*-erg6-gfp* mutant was significantly reduced in the presence of doxycycline, confirming loss of Erg6 at the protein level when *erg6* gene expression was repressed (Fig. 7B, lower right panel). To demonstrate that the punctate localization of Erg6 overlapped with lipid droplets directly, we next stained for lipid droplets in the *erg6-gfp* strain using the lipophilic fluorescent dye BODIPY 558/568 C_12_, a specific tracer of lipid trafficking (29, 30). As detected by fluorescent microscopy, the GFP-labeled puncta overlapped with the red BODIPY staining completely (Fig. 7C). This finding indicated that Erg6-GFP co-localized with lipid droplets in actively growing hyphae. Therefore, our data demonstrate that, similar to *S. cerevisiae*, Erg6 localizes to *A. fumigatus* lipid droplets.

**Figure 7.**
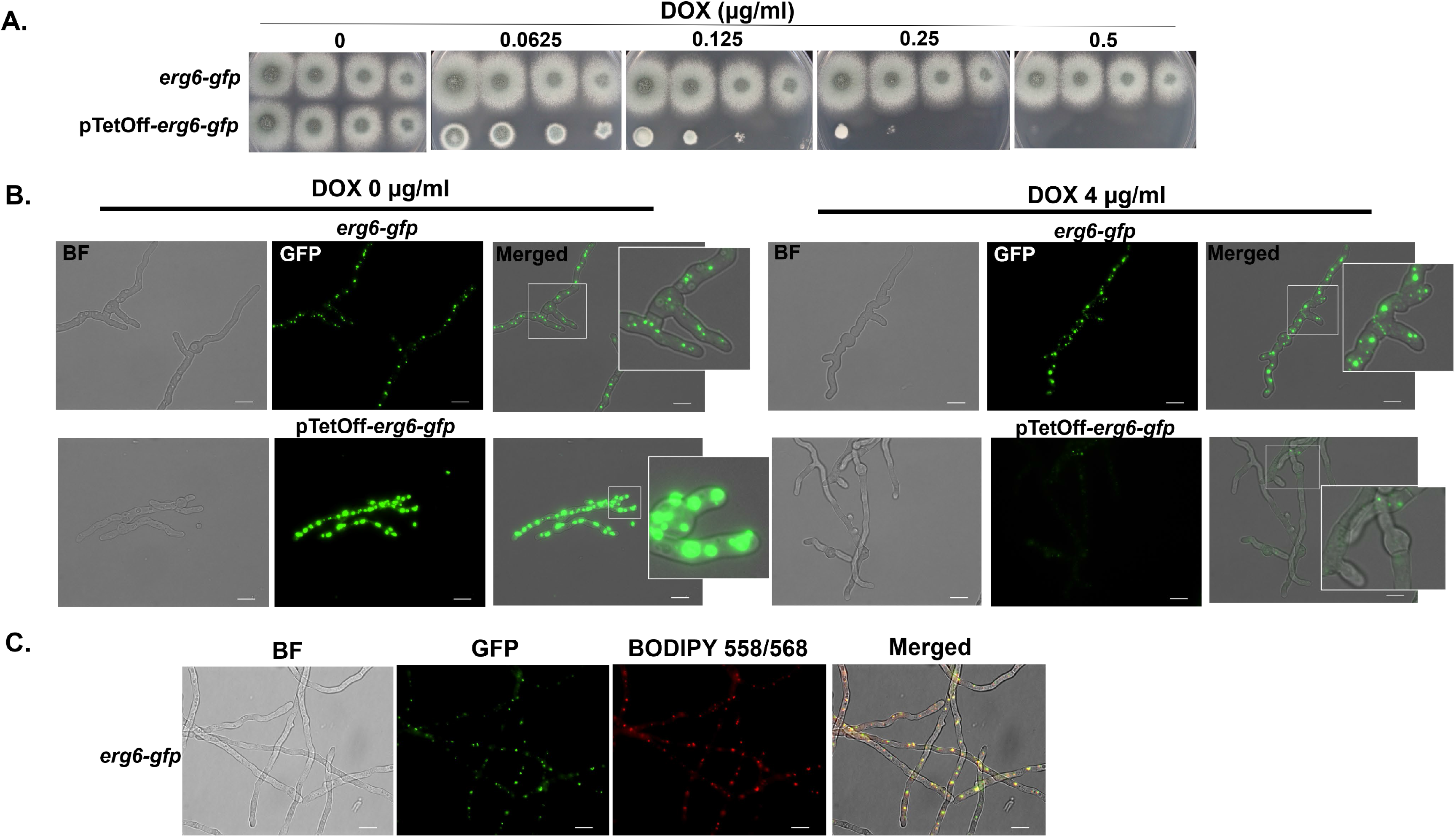
Erg6 localizes to lipid droplets in *A. fumigatus* hyphae. **(A)** Spot-dilution cultures, performed as described in Figure 1B, indicate that fusion of *egfp* to the 3’ end of *erg6* in either the parent or pTetOff-*erg6* background does not negatively affect *erg6* function. Note similarities to growth in untagged strains (Figure 1B). **(B)** Mature mycelia were developed in GMM broth using the indicated doxycycline concentrations for 16 h at 30°C. Fluorescent images were captured using GFP filter settings. **(C)** Co-localization of Erg6-GFP to *A. fumigatus* lipid droplets using the droplet marker, BODIPY 558/568. Conidia of the erg6-gfp strain were culture to mature hyphal development and subsequently stained with 1 µg/ml BODIPY 558/568 C12 for 20 min at room temperature. Images were captured using GFP and TRITC filter settings, respectively. Scale bar=10 µm.

### Repression of *erg6* does not alter susceptibility to ergosterol-targeted antifungals

Multiple classes of currently available antifungal drugs target ergosterol or the ergosterol biosynthesis pathway to destabilize cell membrane integrity and function (31). As *erg6* is required for biosynthesis of ergosterol in yeast species, *erg6* gene mutation has been described as resulting in *Candida* and *Cryptococcus* yeast cells with decreased ergosterol content and increased resistance to the ergosterol-binding antifungal drug, amphotericin B (18, 32). In contrast, *Cryptococcus neoformans erg6* null mutant have been shown to be hypersusceptible to the triazole antifungals, a class of lanosterol-14-a-demethylase inhibitors (18). To explore whether *erg6* repression alters antifungal susceptibility in *A. fumigatus*, we carried out MICs assays in the parental strain and pTetOff-*erg6* mutant by strip-diffusion assays. So that sufficient mycelia were obtained for the pTetOff-*erg6* mutant under repressive conditions to accurately monitor the zone-of-inhibition, we utilized the sub-lethal concentrations of 0.125 and 0.25 μg / ml doxycycline embedded GMM agar plates in combination with voriconazole, isavuconazole, itraconazole, posaconazole, and amphotericin B strips. Unexpectedly, after 48 hours of culture, no significant difference in MIC (2-fold or more change) was noted under any condition (Fig. 8A and 8B). The results of these strip-diffusion tests were consistent with broth micro-dilution antifungal susceptibility testing (Fig. S5). To verify whether the susceptibility profiles under *erg6*-repressed conditions might be affected by other factors, we also measured the expression of two efflux pump genes, *abcC* and *mdr1*, associated with resistance to triazoles (33). Doxycycline treatment had no influence on the expression of either efflux pump in the parental strain (data not shown). Surprisingly, RT-qPCR analysis revealed that *abcC* and *mdr1* were overexpressed 3- to 5-fold (log2) under *erg6* repression conditions compared to the no-doxycycline control (Fig. 8C). Therefore, it is possible that increased efflux pump activity under *erg6* repression might counterbalance the accumulation of antifungals in fungal cells, especially for the triazoles for which efflux is a characterized resistance mechanism.

**Figure 8.**
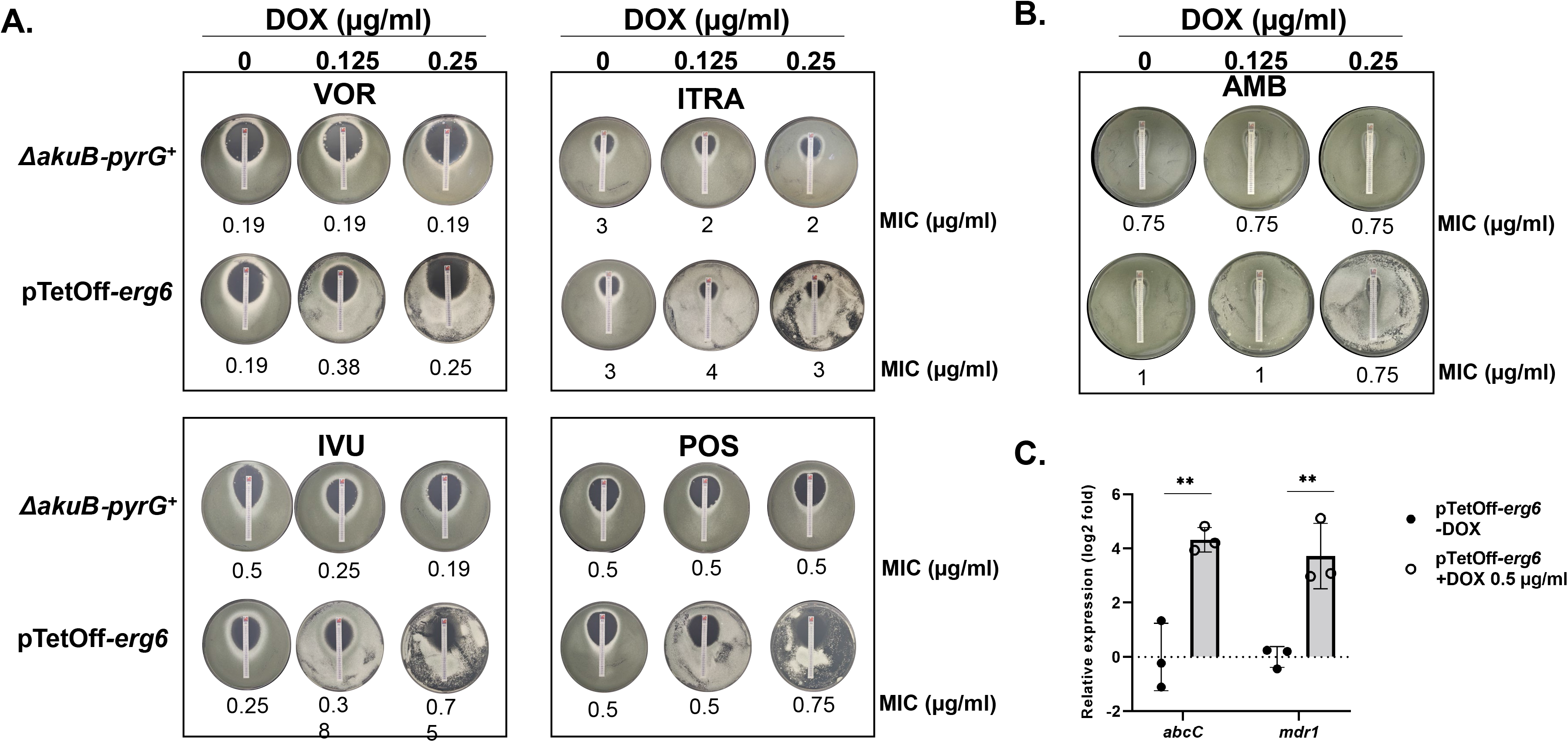
Downregulation of *erg6* does not alter antifungal susceptibility in *A. fumigatus*. Strip-diffusion MIC assays of the parental and TetOff-*erg6* strains were carried out under the indicated doxycycline concentrations. Conidia (2×10^6^) suspended in 0.5 ml sterile water were spread evenly over GMM plates and allowed to dry. Commercial test strips embedded with voriconazole, itraconazole, isavuconazole, posaconazole **(A)** or amphotericin B **(B)** were applied and plates were incubated for 48 h. Resulting MIC values are indicated at the bottom of each plate image. AMB, amphotericin B, VOR, voriconazole, ITRA, itraconazole, IVU, isavuconazole, POS, posaconazole. **(C)**. The expression levels of *abcC* and *mdr1* under the indicated conditions were analyzed by RT-qPCR. Gene expression was normalized to the reference gene, *tubA*, and data is presented as mean ± SD of log_2_ fold change. All assays were performed in biological triplicate. Two-tailed Student t-test was used for data analysis. **p=0.001.

### Repression of *erg6* reduces fungal burden in a murine model of invasive aspergillosis

Our *in vitro* results demonstrate that *erg6* is required for viability of *A. fumigatus* and that addition of exogenous sterols cannot rescue growth. To determine the effect of *erg6* repression *in vivo*, we next compared accumulation of fungal burden during infection with the pTetOff-*erg6* mutant with or without doxycycline in a chemotherapeutically immune suppressed mouse model of invasive aspergillosis. The TetOff system employed here has been validated to regulate expression of a target genes in a murine invasive pulmonary aspergillosis model (34, 35). Previous studies reported that the efficacy of the TetOff system is affected by the starting time of doxycycline administration *in vivo*, and administration beginning 1 day prior to infection yields effective target gene regulation (34). However, it is well documented that intense doxycycline regimens are toxic to mice, resulting in weight loss and lethargy symptoms similar to infection (35). Thus, in this study, we applied doxycycline daily (50 mg/kg, intraperitoneal injection) to the mice beginning 3 days prior to inoculation of conidia. Mice (n=15 per group) were chemotherapeutically immune suppressed with cyclophosphamide and triamcinolone acetonide as described in Materials and Methods and lungs were removed at 40 hrs post-infection. Mice infected with pTetOff-*erg6* displayed a significant reduction of fungal burden in lung tissue under doxycycline treatment compared to that of the no-doxycycline group (*p*=0.0141) (Fig. 9). Therefore, repressing expression of *erg6* interferes with infection progression of invasive aspergillosis *in vivo*.

**Figure 9.**
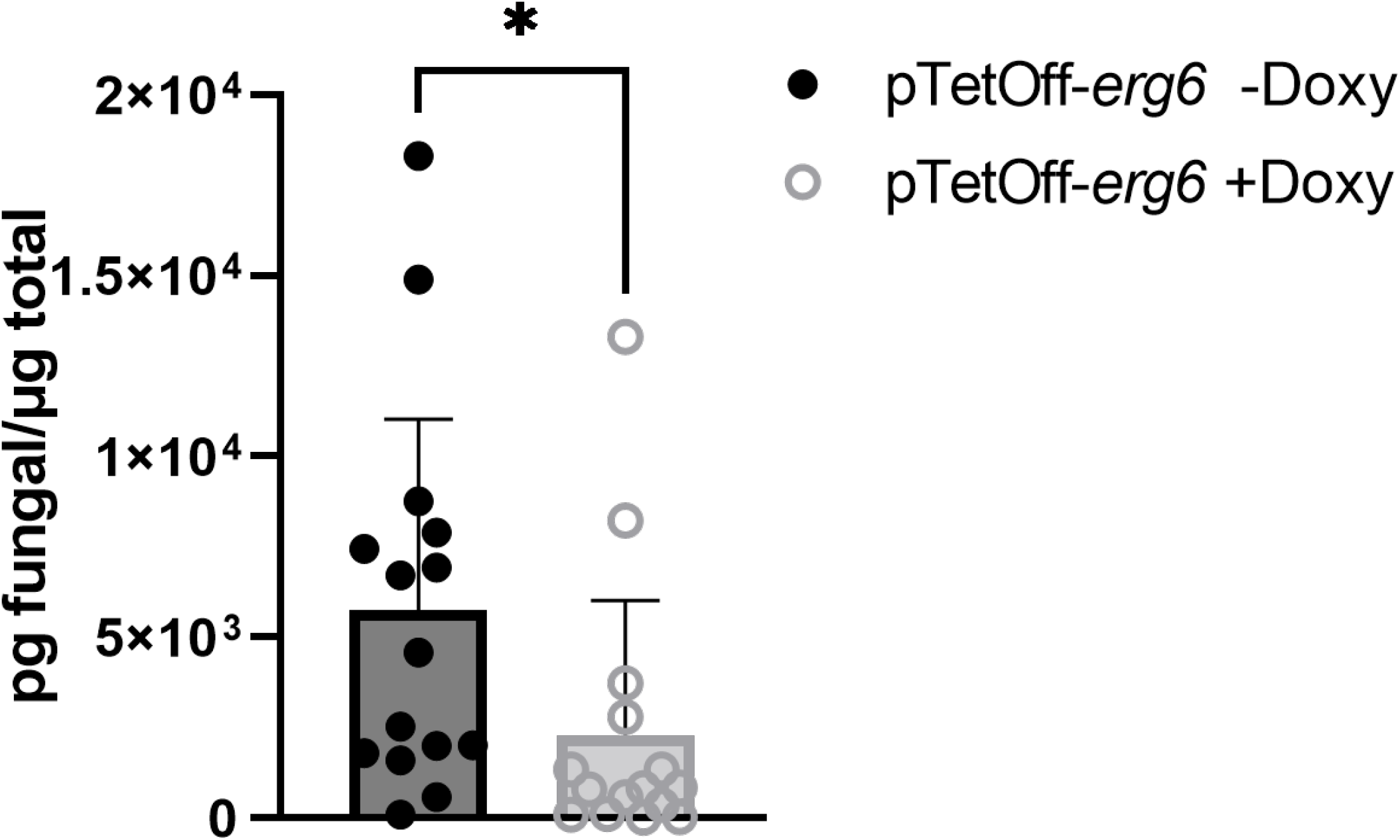
Repression of *erg6* reduces accumulation of fungal burden during *A. fumigatus* infection. Measurement of the lung fungal burden by qPCR at 40 h post infection. Mice (n=15/group) were immune suppressed chemotherapeutically using cyclophosphamide and triamcinolone acetonide as described in Material and Methods and inoculated with 5×10^6^ conidia of the pTetOff-*erg6* strain. For the treatment arm, doxycycline (50 mg/kg) was supplied intraperitoneally once a day beginning on Day −3. Data are represented as pg of *A. fumigatus* DNA in 500 ng of total DNA and is depicted as the mean ± SEM. A Mann-Whitney test was used for statistical analysis. *p<0.02.

## DISCUSSION

Ergosterol is an essential sterol component of the fungal plasma membrane and is involved in numerous architectural and biological functions, such as membrane integrity, fluidity, and permeability (reviewed in (5, 36)). A robust body of literature has demonstrated that disturbed ergosterol homeostasis results in membrane dysfunction and even cell death (37). While ergosterol biosynthesis has been well explored in yeast, studies in filamentous fungi have been relatively limited. In this study, we characterized the sterol C-24 methyltransferase encoding gene, *erg6*, as an essential gene in *A. fumigatus* and analyze roles for Erg6 in growth, ergosterol biosynthesis, drug resistance and establishment of infection.

Some studies have defined *A. fumigatus* having only one copy of sterol C-24 methyltransferase, namely *erg6* (38, 39), whereas others have described *smt1* orthologs as putative *erg6* paralogs (40). Based on our analysis revealing *A. fumigatus* Smt1 to have 31.39% identity to *S. cerevisiae* Erg6 in amino acid sequence, it is possible that *smt1* is a paralog of *erg6* encoding sterol C-24 methyltransferase in *A. fumigatus*. Unlike yeast, it is common for the genomes of filamentous fungi to encode multiple paralogous genes in the ergosterol synthesis pathway. For example, *A. fumigatus* harbors two copies of *erg10*, *erg11*, *erg 24*, *erg25*, and *erg4* while encoding three copies of *erg3* and *erg7* (39). Gene duplications tend to generate functional redundancies and protections against negative effects of genetic mutations (41, 42). The duplicated genes encoding Erg11 (Cyp51), Erg24, Erg25, Erg4, and Erg3 that have been studied thus far in *A. fumigatus* have largely overlapping functions in ergosterol biosynthesis, which allows *A. fumigatus* to maintain partial ergosterol homeostasis under single deletion of these *erg* genes (12, 43–48). In our study, while both Erg6 and Smt1 are characterized as putative sterol C-24 methyltransferases, neither responded to the absence of the other in transcriptional level (Fig. S3C), which is consistent with the observation of two essential paralogs, *erg10A* and *erg10B*, encoding acetyl-CoA acetyltransferase (49). However, our findings mostly support the conclusion that either *smt1* is not a functional paralog of *erg6*, or *smt1* contributes little to C-24 methyltransferase activity in support of ergosterol biosynthesis in *Aspergillus*. For example, the *Δsmt1* mutant demonstrates a wild-type growth phenotype (Fig. S3A), *smt1* overexpression is unable to restore the phenotypic defects resulting from *erg6* deficiency (Fig. 4A), and loss of *smt1* does not cause exaggerated phenotypes in an *erg6* repressed mutant (Fig. 4B). Therefore, if Smt1 functions in ergosterol biosynthesis, we postulate that *erg6* acts as the predominant sterol C-24 methyltransferase and is able to compensate the loss of *smt1*. Further functional experiments still need to be performed to confirm the role of Smt1.

In this study, we validated that lipid droplet-localized Erg6 is essential for *A. fumigatus* survival *in vitro* and is required for fungal infection *in vivo*. As *Δerg6* was unviable, we constructed tetracycline-regulatable *erg6* mutants as an alternative to explore the essentiality of Erg6. Under *erg6* repression, our findings showed that *A. fumigatus* was unable to break dormancy as indicated by conidia showing little metabolic viability (Fig. 2A and 2B). Furthermore, Erg6 is not only required for germination, but also for hyphal growth. When pTetOff-*erg6* mutant was pre-germinated under inducing conditions, germlings were unable to continue growth and support colony development when transferred to repressing culture conditions (Fig. 2C). Given the impaired growth under *erg6* repression *in vitro*, it is not surprising that downregulation of *erg6* caused decreased fungal burden in a chemotherapeutic murine model of aspergillosis (Fig. 9). As we showed that the *erg6* orthologs are essential for *in vitro* survival of *A. lentulus*, *A. terreus* and *A. nidulans* as well (Fig. 3), it is likely that Erg6 activity is essential for pathogenic growth across *Aspergillus* species. The importance of Erg6 to pathogenic fungal fitness appears to be conserved across many fungal pathogens, even though *erg6* orthologs are not essential in most yeast species studied to date. For example, absence of *erg6* does not lead to growth defects in *C. glabrata*, *C. albicans*, or *K. lactis*(*15–17*), and causes only modest growth defects in *C. neoformans* and severe growth defects in *C. lusitaniae* (18, 50). However, a *C. neoformans* Δ*erg6* mutant and mutants with reduced *erg6* expression in *C. albicans* have been reported to have significantly reduced virulent in *G. mellonella* infection models (18, 51). Although *erg6* null mutants in yeast are viable, *erg6* deficiency contributes to compromised phenotypes related to ergosterol-dependent functions, such as increased cell membrane permeability, reduced cell wall integrity, loss of thermotolerance and altered antifungal susceptibility profiles (14–18). Thus, *erg6* may be a promising antifungal drug target.

The outcomes of inhibiting specific steps in the ergosterol biosynthesis pathway are ergosterol deficiency and the accumulation of sterol intermediates. As expected, the substrate of Erg6, lanosterol, is the major accumulated intermediate when *erg6* is repressed, whereas lanosterol is a barely detectable intermediate in non-repressive conditions. It was somewhat surprising that ergosterol remained the most abundant sterol at nearly 45% of the total sterol pool when the pTetOff-*erg6* mutant was grown in the presence of 2 μg / ml doxycycline. Under these conditions on solid agar, mycelial growth was completely inhibited. We attribute this outcome to the differences in attempting doxycycline-regulated gene repression under differing culture conditions (i.e., submerged growth *vs*. sold agar) coupled with the potential leakiness of our TetOff gene regulation system (35). Along with ergosterol and lanosterol dominating the sterol pool, each cholesta-type intermediates (including cholesta-5,7,22,24-tetraenol, cholesta-5,7,24-trienol, 4,4-dimethyl cholesta-dienol and cholesta-dienol) constituted less than 4% of total sterols in pTetOff-*erg6* mutants under doxycycline treatment. These cholesta-type sterols are not detectable in non-repressive conditions (Table 1). These findings differ from the reported sterol accumulation in yeast organisms when Erg6 activity is lost. No detectable ergosterol was measured in the viable *erg6* null mutants of *S. cerevisiae*, *C. albicans*, *C. neoformans*, and *K. lactis* (14, 15, 17, 18). Instead, ergosterol biosynthesis in the absence of *erg6* causes abundant accumulation of zymosterol, cholesta-5,7,24-trienol and cholesta-5,7,22,24-tetraenol. The exact reasons for the differing sterol profiles between *A. fumigatus* and yeast under *erg6* deficiency are still unclear. However, this is likely due to variation in the preferred substrate specificities of Erg6 in different organisms. For example, *S. cerevisiae* Erg6 seems to prefer zymosterol as a substrate whereas lanosterol appears to be the preferred Erg6 substrate in some filamentous fungal organisms (5, 12).

Probably the most explored phenotype related to *erg6* deletion in yeast is the subsequent alteration of antifungal drug susceptibility. Given that the first-line antifungal drugs, triazoles and polyenes, target ergosterol biosynthesis and ergosterol itself, respectively, we hypothesized that the defective ergosterol production resulting from *erg6* repression might affect antifungal susceptibility. Surprisingly, no significant alternations in triazole or polyene resistance profiles were observed in *A. fumigatus* in response to *erg6* repression (Fig. 8). In addition, although we noted that Erg6-GFP localization was maintained in lipid droplets upon triazole stress, *erg6* gene expression was significantly upregulated, (Fig. S6A and S6B). Thus, although *erg6* is transcriptionally responsive to triazole-mediated pathway perturbation, loss of Erg6 activity does not appear to synergize with triazole therapy. The wild type resistance profiles we observed under *erg6* repression are in stark contrast to reports from *S. cerevisiae*, *K. lactis*, *C. neoformans*, *C. albicans*, and *C. glabrata* in which *erg6* deletion is associated with increased resistance to polyenes (15–18, 52). This acquired polyene resistance is thought to be underpinned by ergosterol reduction or depletion in these mutants. The resistance is then directly related to the mechanism of action of polyene drugs which is to bind ergosterol, and extract it out of the cellular membrane to cause lethality (53). Therefore, high MICs to polyenes are commonly seen in ergosterol-defective strains (54). As for triazoles, the susceptibility profiles are species- and drug-dependent. Increased susceptibility has been reported for *erg6* null mutants of *S. cerevisiae*, *K. lactis* and *C. neoformans* (15, 18), whereas a C. *albicans Δerg6* mutant maintains wild-type susceptibility profiles (17). *C. glabrata* clinical isolates with *erg6* mutation displays increased susceptibility to triazoles (32), but null mutants generated in laboratory strains revealed increased tolerance (16). Additionally, reduced resistance to triazoles has been reported in several *erg* null mutants along with ergosterol reduction or depletion (44, 46, 48). The mechanisms involved in alternation of triazole susceptibility caused by *erg6* mutation are complicated. The disturbed membrane fluidity and permeability caused by altered ergosterol biosynthesis, which allows azoles to penetrate abnormally, is viewed as the main potential explanation (55). One possible explanation for the lack of changes in antifungal drug resistance profiles between wild type and *erg6* repressed strains in this study is that total cellular ergosterol content under the levels of *erg6* repression obtained here may still be sufficient to maintain the plasma membrane functionally. It should be noted that the highest doxycycline level we used in our MIC assays was 0.25 µg / ml, which allowed observable mycelia to grow in the plates so that we could reliably measure the zone-of-inhibition. Additionally, we found that *erg6* repression triggered the overexpression of two triazole resistance-associated efflux pump genes, *abcC* and *mdr1* (Fig. 8C). Therefore, increased efflux could theoretically abrogate any increased sensitivity to triazoles that may have resulted from ergosterol depletion in *erg6*-repressed strains. Further analyses of efflux pump activity changes in response to *erg6* repression are needed.

In conclusion, we have validated *A. fumigatus* Erg6 as an essential protein that localizes to lipid droplets and regulates ergosterol biosynthesis. We also found that Erg6 orthologs are essential for viability in additional *Aspergillus* species *in vitro* and that *A. fumigatus* Erg6 is required for establishment of fulminant infection and invasive aspergillosis mouse model. Given the importance of Erg6 for growth, virulence and drug susceptibility patterns across fungal pathogens, our data support Erg6 as a promising target for antifungal drug development.

## MATERIALS AND METHODS

### Strains and growth conditions

All strains used in this study are summarized in Table S1. All strains were routinely cultured at 37°C on Glucose Minimal Medium (GMM) agar plates, supplemented with 5% yeast extract, as necessary (56). Conidia were harvested from GMM plates using sterile water and stored at 4°C. For spot dilution assays, GMM agar plates containing doxycycline at the indicated concentrations were point-inoculated with serial dilutions of conidial suspensions from 50,000 to 50 conidia. The plates were incubated at 37°C for 48 h. Hyphal morphology of submerged culture was assessed by inoculating 10^6^ conidia into the wells of 6-well plates containing liquid GMM at the indicated doxycycline concentrations and sterile coverslips. After 16 h at 37°C, coverslips were washed twice with PBS and mounted for microscopy. For post-germination growth assays, 10^7^ conidia were cultured in 10 ml GMM broth for 8 h. After confirming germling formation by microscopy, ten microliters of germling suspension were inoculated onto fresh GMM plates containing doxycycline at the indicated concentrations for sub-culture for 48 h at 37°C.

### Construction of mutant strains

Genetic manipulations in this study were performed using a CRISPR-Cas9 gene editing techniques described previously (20). Briefly, for CRISPR-Cas9-mediated gene deletion, two PAM sites located upstream and downstream of the desired genes were selected and used for crRNA design. Repair templates, composed of a hygromycin resistance cassette, were amplified using primers flanked with 40 bp microhomology regions of the target locus (Table S2). For overexpression mutants, native promoters of target genes were replaced by the *hspA* promoter (20) by identifying and utilizing a single PAM site upstream of the gene coding region, as we have previously described (20). Similarly, doxycycline-regulatable *erg6* mutants were generated via pTet-Off and pTetOn promoters, as previously described (21, 22). Ribonucleoprotein (RNP) complexes were assembled *in vitro* using commercially available crRNA, tracrRNA, and the Cas9 enzyme as described previously (57). Briefly, equal molar amounts of crRNA and tracrRNA were mixed in duplex buffer and boiling at 95°C for 5 min. After cooling at room temperature for 10 min, duplex crRNA-tracrRNA was combined with Cas9 enzyme (1μg / μl), followed by incubation for 5 min at room temperature. Transformation was performed as described previously (57). Transformation mixtures containing 10 μl protoplasts (1 − 5×10^5^ cells), 5 μl RNP (described above), repair template (900 ng), 3 μl polyethylene glycol (PEG)-CaCl_2_ buffer and STC buffer (1.2 M sorbitol, 7.55 mM CaCl_2_·H_2_O, 10 mM Tris-HCl, pH 7.5) were incubated on ice for 50 min. Subsequently, the mixture was added to 57 μl polyethylene glycol (PEG)-CaCl_2_ buffer and incubated at room temperature for 20 min. The mixture was brought to 200 μl STC buffer and plated onto Sorbitol Minimal Medium (SMM) agar plate finally. After overnight room temperature incubation, transformation plates were overlaid with SMM top agar containing selective drug and incubated at 37 °C until colonies were observed. For the generation of Erg6-GFP strain, a repair template was amplified using a GFP-expression vector that contained a linker sequence (AGATCTGGATGCGGCCGC) flanked with 40 bp microhomology regions at the 3’ end of *erg6* (excluding the termination codon) to direct integration at a single downstream PAM site. All mutants were confirmed by multiple genotyping PCR reactions to ensure proper integration of the introduced repair template. All PAM sites and protospacer sequences used for crRNA design are included in Table S2.

### RNA extraction and quantitative real-time PCR analysis

RNA extraction and RT-qPCR were carried out as previously described (22). In brief, all strains were cultivated in liquid GMM supplemented with 5% yeast extract at 37°C/250 rpm for 18 h. Mycelia were harvested, frozen in liquid nitrogen, and ground using a pestle and mortar. Total RNA was extracted using Qiagen RNeasy Mini Kit following the manufacturer’s protocol. DNA contamination from RNA samples was eliminated by RNase-free Turbo DNase Kit (Invitrogen). Subsequently, cDNA was synthesized using SuperScript II system (Invitrogen), following the manufacturer’s instructions. Quantitative real-time PCR was carried out using SYBR® Green Master Mix (Bio-Rad) in a CFX Connect Real-Time System (Bio-Rad).

### Antifungal susceptibility assay

The susceptibility profiles of antifungals including amphotericin B, itraconazole, voriconazole, Posaconazole, and isavuconazole were evaluated using commercial E-test strips following the manufacturer’s protocol. Briefly, 2×10^6^ conidia in 0.5 ml were spread onto GMM plates containing the indicated doxycycline concentrations. The antifungal embedded strips were applied onto the dried agar plates. After 48 h of culture, the MICs were measured by observation of zone-of-clearance.

### Fluorescence microscopy

To visualize Erg6-GFP localization, approximately 10^6^ conidia were cultured in liquid GMM on sterile coverslips at indicated concentrations of doxycycline or antifungal drugs for 16 h at 30°C or 37°C. For Live/Dead staining, coverslips were washed once with 0.1M MOPS buffer (pH 3) and stained with 50 μg / ml 5,(6)-Carboxyfluorescein Diacetate (CFDA) (Invitrogen) in MOPS buffer for 1 h at 37°C in the dark. For lipid droplet staining, coverslips were stained with 1 µg / ml BODIPY 558/568 C_12_ in PBS buffer for 30 min at room temperature. For ergosterol staining, hyphae cultured on coverslips were stained with filipin (Sigma) at the final concentration of 25 μg / ml in liquid GMM for 5 min. After the above staining procedures, coverslips were washed twice with indicated buffer and mounted for microscope. Fluorescence microscopy was performed on a Nikon NiU microscope. CFDA and GFP staining were visualized using GFP filter settings. Lipid droplet fluorescence was captured using TRITC filter settings. Filipin staining was observed using DAPI filter settings. Images were captured by Nikon Elements software (version 4.60).

### Sterol extraction and composition analysis

Conidia were cultured in RPMI-1640 medium buffered with 0.165 M MOPS (pH 7.0) containing 0.2% w/v glucose at a final concentration of 1×10^6^ cells / ml in the indicated concentrations of doxycycline for 16 h at 37°C/250 rpm. Mycelia were harvested and non-saponifiable lipids were extracted as previously described (58). Briefly, sterols were derivatized using 0.1mL BSTFA TMCS (99:1) and 0.3 mL anhydrous pyridine and heating at 80°C for 2 h. TMS-derivatized sterols were analyzed using GC/MS (Thermo 1300 GC coupled to a Thermo ISQ mass spectrometer, Thermo Scientific) and identified with reference to relative retention times, mass ions, and fragmentation spectra. GC/MS data files were analyzed using Xcalibur software (Thermo Scientific). Sterol composition was calculated from peak areas, as a mean of three replicates. Data was presented as mean percentage ± SD for each sterol.

### Murine model of invasive pulmonary aspergillosis

All animal studies were performed under the guidance of the University of Tennessee Health Science Center Laboratory Animal Care Unit and approved by the Institutional Animal Care and Use Committee. Animal models of infection were performed as previously described (59). CD-1 female mice (Charles River or Envigo) weighing approximately 25 g were chemotherapeuticaly immune suppressed by intraperitoneal injection of 150 mg / kg of cyclophosphamide (Sigma-Aldrich) on days −3, +1, +4 and +7, and subcutaneous injection of 40 mg/kg triamcinolone acetonide (Kenalog, Bristol-Myers Squibb) on day −1. Doxycycline was supplied at 50 mg/kg by intraperitoneal injection once per day from day −3. One day 0, mice were anesthetized with 5% isoflurane and intranasally infected with a dose of 5×10^6^ conidia in 40 μl saline solution. After 40 h of infection, mice were humanely euthanized by anoxia with CO_2_. Lungs were harvested and immediately frozen in liquid nitrogen. Frozen tissue was lyophilized (SP Scientific VirTis Benchtop Pro BTP) for 48 h and subsequently ground to fine powder by using a beat beater (Bullet Blender Gold). Fungal DNA extraction was performed following previously described protocols (60). qPCR analyses were conducted by PrimeTime Gene Expression Master Mix and qPCR Probe Assays (Integrated DNA Technologies) with primers to amplify the *A. fumigatus* 18S rRNA gene as previously described (61). To measure the fungal burden, a standard curve containing five dilutions from 100 to 0.01 ng of *A. fumigatus* genomic DNA (10-fold dilution) was calculated to determine the amount of fungal DNA. For each sample, 500 ng of total DNA from pulverized lung tissue was used as template in qPCR assay. A negative control, using H_2_O to replace the DNA template, was employed. The qPCR protocol was run on a CFX Connect Real-Time System (Bio- Rad). Technical triplicates were conducted for each lung sample. The fungal burden was calculated as nanograms of *A. fumigatus* specific DNA in 500 ng of total DNA.

## FIGURE LEGENDS

**Supplemental Figure 1. Phylogenetic analysis of the homologs of sterol C-24 methyltransferase encoding genes from selected model and pathogenic fungi.** The putative full-length amino acid sequence of each organism was acquired by BLASTP analysis using the *erg6* of *S. cerevisiae* (SGD: S000004467) as a query sequence against the indicated database. Alignment analysis was performed by CLUSTALW and a phylogenetic tree was constructed by MEGA 11 software using the maximum likelihood method with a bootstrap value of 1000. Organisms used for comparison are *A. fumigatus*, *A. lentulus*, *A. terreus*, *A. nidulans*, *Neurospora crassa*, *S. cerevisiae, C. albicans* and *C. neoformans*.

**Supplemental Figure 2. Schematic of gene manipulations by CRISPR/Cas9 editing. (A)** Deletion of the gene of interest (GOI). Two protospacer adjacent motifs (PAMs, indicated as red bars) flanking the GOI were targeted by repair templates composed of a hygromycin resistance cassette with ~40-basepair microhomology regions for integration upstream and downstream of the GOI in the Δ*akuB*-pyrG+ genetic background to generate Δ*smt1* and Δ*erg6* mutant. **(B)** Tetracycline repressible or inducible expression and overexpression of GOI. A PAM site upstream of the GOI was targeted with a repair template carrying the TetOff or TetOn promoter construct or the strong pHspA promoter fused to either a phleomycin or hygromycin resistance cassette. **(C)** Generation of GFP-tagged Erg6. A repair template containing a linker sequence, the egfp coding sequence, and a phleomycin resistance cassette was amplified using primers to incorporate microhomology regions on either side of a PAM stie selected at the 3’ end of *erg6*.

**Supplemental Figure 3. Analyses of the putative *A. fumigatus erg6* paralog, *smt1*. (A)** Colony morphology of the parental strain, Δ*smt1*, and OE-*smt1*. A total of 10,000 conidia were inoculated onto GMM agar plates and incubated for 48 h at 37°C. **(B)** The expression level, as measured by RT-qPCR, of *smt1* after promoter replacement mutation using the pHspA promoter. **(C)** Expression changes in *smt1* and *erg6* in response to loss of the respective paralogs. Mycelia were harvested after 16 h in liquid GMM at 37°C/250 rpm. Gene expression was normalized to the reference gene, *tubA*, and data presented as mean ± SD of log_2_ fold change. All assays were performed in biological triplicate. Two-tailed Student t-test was used for statistical analysis. ***p=0.0005, ****p<0.0001.

**Supplemental Figure 4. Growth of the pTetOn-*erg6* mutant is doxycycline dependent.** Spot-dilution assays of the parental and pTetOn-*erg6* strains were performed on GMM agar plates using the indicated doxycycline levels. Culture conditions were as described in Figure 1B.

**Supplemental Figure 5. Repression of *erg6* does not alter antifungal susceptibility profiles in *A. fumigatus*.** Broth dilution antifungal susceptibility assays were performed in triplicate for each strain using the indicated doxycycline concentration. Assays were conducted according to the CLSI standard M38-A2.

**Supplemental Figure 6. Voriconazole treatment increases *erg6* expression but does not alter protein localization. (A)** RT-qPCR analysis of *erg6* expression with or without voriconazole treatment. Mycelia were harvested after 16 h in liquid GMM supplemented with or without 0.125 µg/ml voriconazole at 37°C/250 rpm. Gene expression was normalized to the reference gene, *tubA*, and data is presented as mean ± SD of log_2_ fold change. All assays were performed in biological triplicate. Two-tailed Student t-test was used for statistical analysis. **p=0.001. **(B)** Mycelia were cultured in GMM broth with 0.125 µg/ml voriconazole for 16 h at 37°C. Fluorescent images were captured using GFP filter settings. Scale bar=10 µm.

## ACKNOWLEDGEMENTS

This work was supported by National Institutes of Health (NIH) / National Institute of Allergy and Infectious Diseases (NIAID) grants R01 AI158442 (JRF) and R01 AI143197 (JRF/PDR). The authors would also like to thank Nathan P. Wiederhold, PharmD at the Fungus Testing laboratory for providing the *A. lentulus* isolate used in this study.

