## Supplemental Figures and Tables for "The sterol C-24 methyltransferase encoding gene, *erg6*, is essential for viability of *Aspergillus* species"

### Supplemental Figure 1

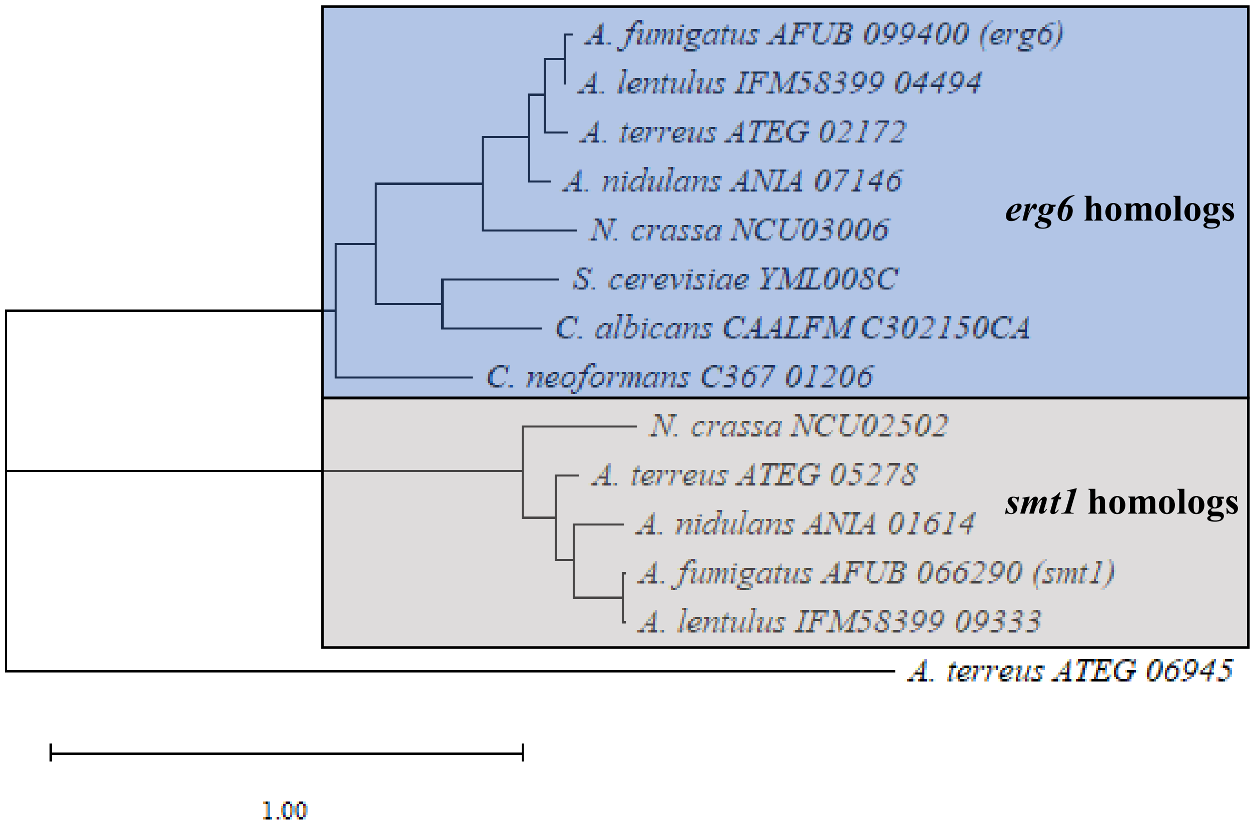

**Supplemental Figure 2**

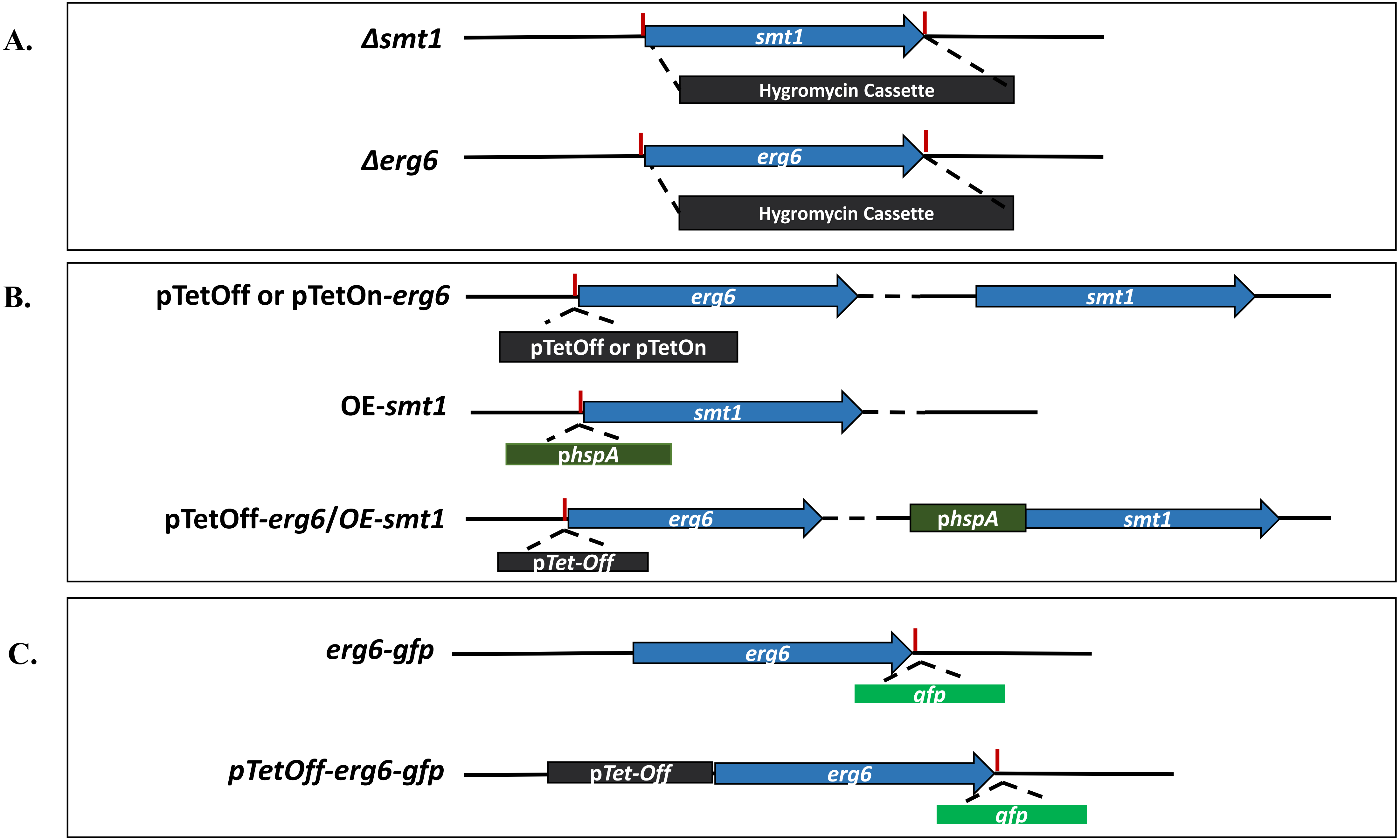

#### Supplemental Figure 3

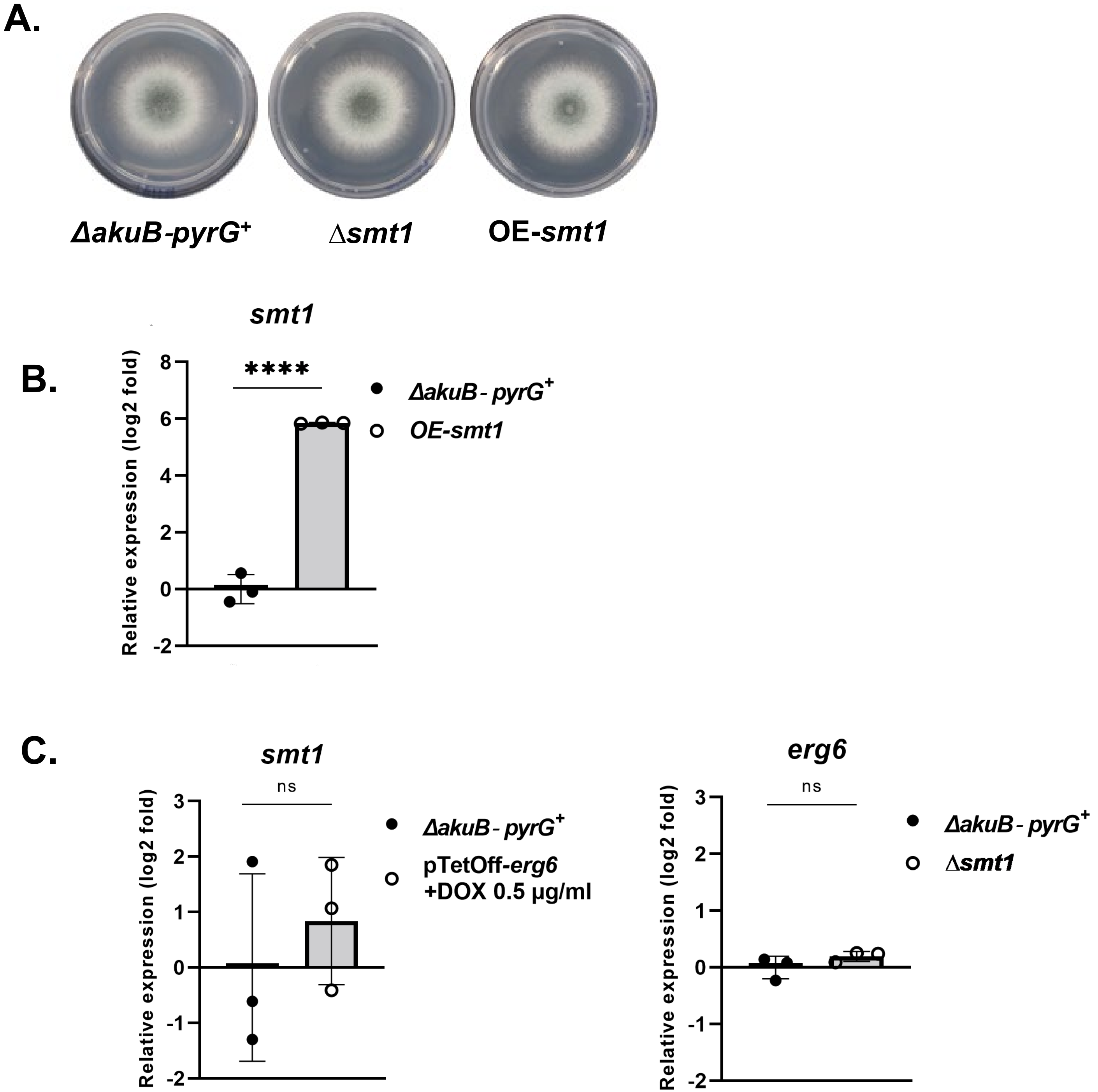

### Supplemental Figure 4

DOX (μg/ml)

0 20 60 100 200

*ΔakuB-pyrG*⁺

pTetOn-*erg6*

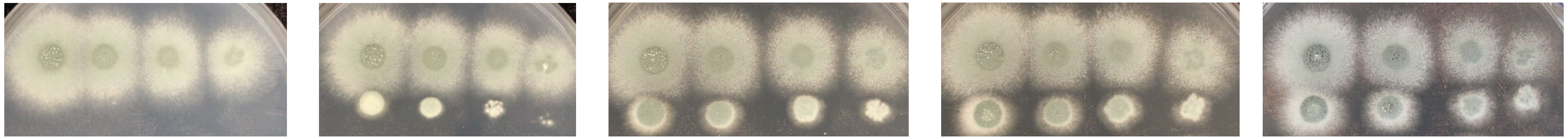

### Supplemental Figure 5

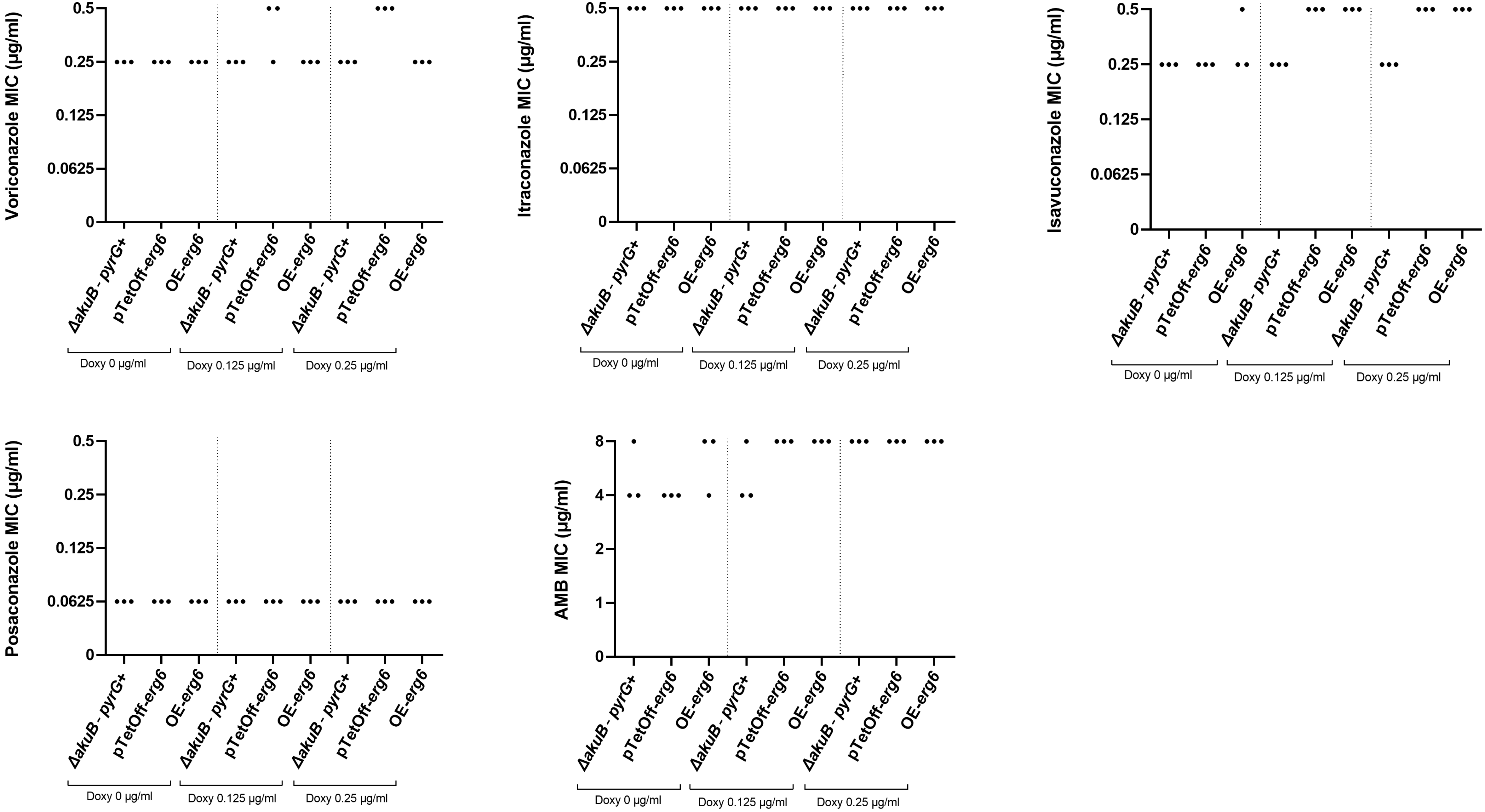

### Supplemental Figure 6

**A.**

***erg6***

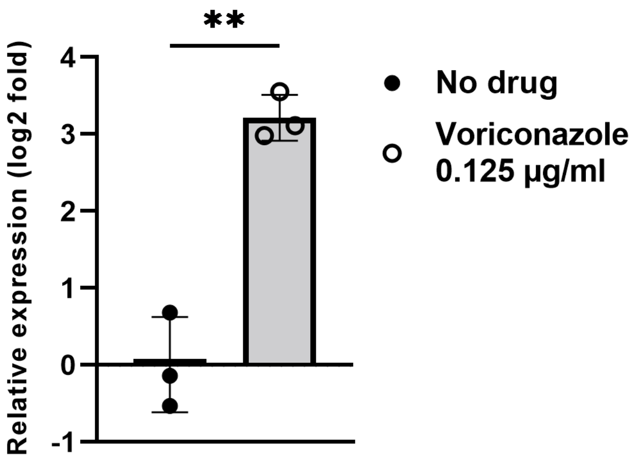


**B.**

***erg6-gfp***

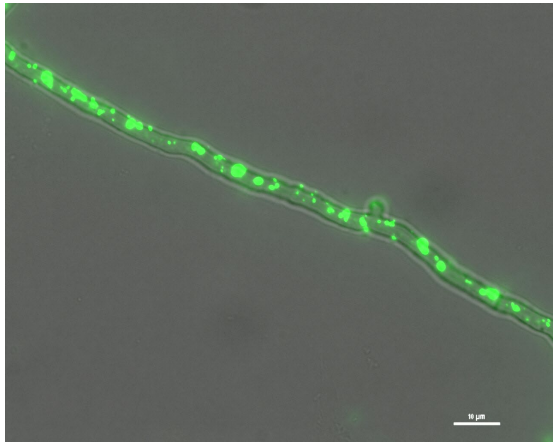

**Voriconazole 0.125 µg/ml**

**Supplemental Table 1. Strains used in this study**

| Species | Strain | Genetic background | Source |
|---|---|---|---|
| *A. fumigatus* | Δ*akuB*-*pyrG*⁺ | KU80Δ*pyrG* | (1) |
| | Δ*smt1* | Δ*akuB*-*pyrG*⁺ | This study |
| | pTetOff-*erg6* | Δ*akuB*-*pyrG*⁺ | This study |
| | pTetOn-*erg6* | Δ*akuB*-*pyrG*⁺ | This study |
| | OE-*smt1* | Δ*akuB*-*pyrG*⁺ | This study |
| | pTetOff-*erg6 /* OE-*smt1* | OE-*smt1* | This study |
| | pTetOff-*erg6 /* Δ*smt1* | Δ*smt1* | This study |
| | *erg6*-gfp | Δ*akuB*-*pyrG*⁺ | This study |
| | pTetOff-*erg6*-gfp | pTetOff-*erg6* | This study |
| *A. lentulus* | DI-19-116 | - | Gift from Nathan P. Wiederhold, PharmD |
| | pTetOff-erg6 / DI-19-116 | DI-19-116 | This study |
| *A. terreus* | FGSC A1156 | - | Fungal Genetics Stock Center |
| | pTetOff-*erg6* / FGSC A1156 | FGSC A1156 | This study |
| *A. nidulans* | FGSC A1166 | - | Fungal Genetics Stock Center |
| | pTetOff-*erg6* / FGSC A1166 | FGSC A1166 | This study |

**Supplemental Table 2. Primers in this study**

| **Purpose** | **Primer name** | **Sequence** |
|---|---|---|
| Protospacer Sequence | Af_Cr_erg6_5’ | AAGTCCAATTGCTATCGCCA |
| Protospacer Sequence | Af_Cr_erg6_3’ | TGTAAAAGAGACTCGTTACT |
| Protospacer Sequence | Af_Cr_smt1_5’ | TGTGGGCGGTGATAGCGGAC |
| Protospacer Sequence | Af_Cr_smt1_3’ | TCATTGCTCAAAAGCCTTCG |
| Protospacer Sequence | At_Cr_erg6_5’ | ACCGCCACCCGTTCTCGCCA |
| Protospacer Sequence | An_Cr_erg6_5’ | ACTGTCAATTCAATTCACAA |
| Protospacer Sequence | Al_Cr_erg6_5’ | AAGTCCAATTGCTATCGCCA |

| | | |
|---|---|---|
| Repair Template | Af_erg6_del RT-F | GATCTGTGATCCACCCCTTTTCCACCCTTACCACATCCAAAGCTTGCATGCCTGCAGG |
| Repair Template | Af_erg6_del RT-R | AGGATGCAGGCACAGGGCAGATATTTGTACAGGCAATTCGCCGAGCTCCCAAATCTGTCCAG |
| Repair Template | Af_erg6_OE RT-F | CTGATCTGTGATCCACCCCTTTTCCACCCTTACCACATCCATTCGTTGACCTAGCTGATTCTGG |
| Repair Template | Af_erg6_OE RT-R | GCGCAAGTGGTTCTCTTGTTCCAAAGCTACGGGGGCCATGGGGATCGAATTCCTGCAGCC |
| Repair Template | Af_smt1_del RT-F | AGCATATCTCAACCTGCCTTGGAATCCAGCTTCTCTTTCTAGCTTGCATGCCTGCAGG |
| Repair Template | Af_smt1_del RT-R | CCAATTGCTCTTGTTTTTTGAGCGGGTGGTTAACAACTGACCGAGCTCCCAAATCTGTCCAG |
| Repair Template | Af_smt1_OE RT-F | AGCATATCTCAACCTGCCTTGGAATCCAGCTTCTCTTTCTAGCTTGCATGCCTGCAGG |
| Repair Template | Af_smt1_OE RT-R | GCCAGGGCCGGTGCTGTGGTCTGTGTTTCCATCATTGTTGTGTGAAGAAGTGAGGAGGGTTTCGT |
| Repair Template | Af_erg6_TetOff RT-F | GATCTGTGATCCACCCCTTTTCCACCCTTACCACATCCAACAATTAAAGCCTTCGAGCGTCC |
| Repair Template | Af_erg6_TetOff RT-R | CGCGCAAGTGGTTCTCTTGTTCCAAAGCTACGGGGGCCATCCGGTGATGTCTGCTCAAGC |
| Repair Template | At_erg6_TetOff RT-F | AATTCCGTCCACCCTCATTCCTCCCCCTTTTTTTACCCAGAATTAAAGCCTTCGAGCGTCCC |
| Repair Template | At_erg6_TetOff RT-R | CACGAGAGTGGTCCTCGCGTTCGAGAGCGGTGGGAGCCATCCGGTGATGTCTGCTCAAGC |
| Repair Template | An_erg6_TetOff RT-F | CTTTCGCCTCCCCCCTTTTCAACCTCCTACTTTTCGATTCAATTAAAGCCTTCGAGCGTCCC |
| Repair Template | An_erg6_TetOff RT-R | CGCGCTGGTGGTTCTCCTTCTCTAAAGCAGTGGGAGCCATCCGGTGATGTCTGCTCAAGC |
| Repair Template | Al_erg6_TetOff RT-F | GATCTGTGATCCACCCCTTTTCCACCCTTACCACGTCCAAAATTAAAGCCTTCGAGCGTCCC |
| Repair Template | Al_erg6_TetOff RT-R | CGCGCGCGTGATTCTCCTGTTCCAAAGTAGCAGGGGCCATCCGGTGATGTCTGCTCAAGC |
| Repair Template | Af_erg6_TetOn RT-F | GATCTGTGATCCACCCCTTTTCCACCCTTACCACATCCAAAATTAAAGCCTTCGAGCGTCCC |
| Repair Template | Af_erg6_TetOn RT-R | CGCGCAAGTGGTTCTCTTGTTCCAAAGCTACGGGGGCCATGTGATGTCTGCTCAAGCGGG |
| Repair Template | Af_erg6_GFP RT-F | CTTCACGCCCATGTATTTGATGGTCGGACGCAAGCCCGAGAGATCTGGATGCGGCCGCATGGTGAGCAAGGGCGAGG A |
| Repair Template | Af_erg6_GFP RT-R | GGCAGATATTTGTACAGGCAATTCGTGTAAAAGAGACTCGAGCTTGCATGCCTGCAGG |
| PCR screening | Af_erg6_in5-F | GGCAGAATGGTCAGGTACTGC |
| PCR screening | Af_erg6_in5-R | CAATGTGGCATACTCAGCACGG |
| PCR screening | Af_erg6_in3-F | GCCGGATCGTTCAAGCACATG |
| PCR screening | Af_erg6_in3-R | GCCAATTACATAGGCAGGTCTCATG |

| | | |
|---|---|---|
| PCR screening | Af_smt1_in-F | ACCCATGGCAGGTCATATGGC |
| PCR screening | Af_smt1_in-R | CTCGTCCGACCAGTCAAGGT |
| PCR screening | Af_erg6_det-F | GGAACAAGAGAACCACTTGCGC |
| PCR screening | Af_erg6_det-R | CGTGAAGAGCTTCTTCTCACCACC |
| PCR screening | Af_smt1_det-F | AGCTCCGTGGACGTCATTG |
| PCR screening | Af_smt1_det-R | TGCGGTAACTCGTACTGCAGAG |
| PCR screening | Af_erg6_del-F | ACTTCTGTCAGAACTCGCAGTTCC |
| PCR screening | Af_erg6_del-R | CCATCAGTGCATACCCAGCAG |
| PCR screening | Af_smt1_del-F | GCAGGTCATATGGCGATTATCCG |
| PCR screening | Af_smt1_del-R | CTGTGAGTCGTGACACGTGC |
| PCR screening | At_erg6_in-F | CGTCTCAATTGCTTGCTGCC |
| PCR screening | At_erg6_in-R | AGCCGACATCCAGCACCTTC |
| PCR screening | An_erg6_in-F | GCGGTGTCGTGATTGTACAATCG |
| PCR screening | An_erg6_in-R | CCAACATCGAGCACCTTCATGC |
| PCR screening | Al_erg6_in-F | TGCTTCAAGATTGTGATCCTGTGG |
| PCR screening | Al_erg6_in-R | TGGTGAGCCAGGTAGTGTTCATG |
| qRT-PCR | Af_erg6-qPCR-F | ACCCGACACTATTACAACCTGGC |
| qRT-PCR | Af_erg6-qPCR-R | GGTGAGCCAGGTAGTGTTCATGAC |
| qRT-PCR | Af_smt1-qPCR-F | TCTGGTGAAGGTTCACGCACTG |
| qRT-PCR | Af_smt1-qPCR-R | GAGCAATGATGCGGTAACTCGTAC |
| qRT-PCR | Af_abcC-qPCR-F | CTGGAGAAGGTCTCAATGTCGAAC |
| qRT-PCR | Af_abcC-qPCR-R | TTGGCCGTGCTTGGTAAGAG |
| qRT-PCR | Af_mdr1-qPCR-F | TCGTTATGTCACTCCCTGAGGG |
| qRT-PCR | Af_mdr1-qPCR-R | GATTCGGAGTCAAGAGCAGATGTG |
